# Periodic and aperiodic contributions to EEG delta power are translatable and complementary Angelman syndrome biomarkers

**DOI:** 10.1101/2025.09.08.674941

**Authors:** Daniel P. Montgomery, Jeremy J. Shide, Adam M. Didouchevski, Abigail H. Dickinson, Michael S. Sidorov

## Abstract

Angelman syndrome (AS) is a neurodevelopmental disorder caused by loss of maternal *UBE3A* expression. With promising AS therapies now in clinical trials, there is a pressing need for reliable and translatable biomarkers. Elevated EEG delta power is a hallmark of AS and a promising biomarker, but traditional measures conflate delta oscillations with broadband spectral shifts, limiting interpretability and utility. We dissociated periodic and aperiodic contributions to delta power using spectral parameterization in children with AS and *Ube3a* mutant mice. In both species, elevated delta power reflected a combination of increased periodic delta oscillations and elevated aperiodic slope and offset. These features were linked to different behavioral domains and followed divergent developmental trajectories, suggesting distinct underlying mechanisms. Together, our findings establish aperiodic changes as a novel translatable EEG biomarker for AS and support the complementary use of periodic and aperiodic features in preclinical and clinical research.

## Introduction

Angelman syndrome (AS) is a neurodevelopmental disorder characterized by speech and motor impairments, seizures, intellectual disability, and abnormal sleep^1,2^. AS is caused by the loss of function of the maternal copy of the *UBE3A* gene encoding an E3 ubiquitin ligase^3,4^. In neurons, the paternal allele of *UBE3A* is epigenetically silenced^5,6^, leaving the maternal copy as the sole source of *UBE3A* expression. This neuron-specific imprinting has made unsilencing the paternal allele a promising therapeutic strategy^7–10^, alongside efforts to restore UBE3A expression more broadly or target downstream pathways pharmacologically^9^. Several potential treatments have advanced to Phase 3 clinical trials (NCT06914609, NCT06617429), marking a major step towards targeted treatment for AS. As efforts continue across both preclinical and clinical development, there is a growing need for objective, reliable, and quantifiable biomarkers to assess treatment efficacy.

Electroencephalography (EEG) has emerged as a promising candidate for biomarker development in AS. Clinically observable increases in delta (∼2-4 Hz) oscillations are present in ∼80-90% of EEGs in children with AS^11^, and are highly specific to AS relative to other neurodevelopmental disorders^12,13^. Delta oscillations are quantifiable through spectral analysis, which enables objective measurement of oscillatory activity across defined frequency bands. Using these methods, elevated delta power in AS has been consistently observed across the scalp, during both wakefulness and sleep, and in children across a wide age range^13–16^. Importantly, elevated delta power correlates with the severity of cognitive impairment in AS^17,18^, suggesting that it may serve as a sensitive, quantifiable measure for evaluating treatment response in clinical trials. Indeed, delta power demonstrated responsiveness to treatment in a recent Phase 1/2 clinical trial^19^.

The utility of delta power as an AS biomarker is further strengthened by its strong cross-species translatability. Rodent models of AS, generated by knocking out the maternal allele of *Ube3a (Ube3a^m-/p+^*), exhibit behavioral impairments with strong face validity across multiple behavioral domains^20^ and show elevated delta power in local field potential (LFP) recordings^14,21–24^. Delta power has also been used as a readout in preclinical studies testing genetic and pharmacological interventions for AS^25–27^. However, the delta phenotype reported in mice has been somewhat less consistent than in humans [e.g., see ^28^]. Moreover, it is unclear whether the strength of delta oscillations meaningfully predicts behavioral impairments in mice, limiting its utility for preclinical research.

Compounding these challenges, both *Ube3a^m-/p+^* mice and children with AS exhibit a general increase in broadband EEG power not limited to the delta range^14,15,22,24,25^, complicating attempts to isolate and quantify delta oscillations above this elevated background. As a result, using raw delta power as a biomarker can conflate true oscillatory activity with nonspecific increases in overall power, limiting its interpretability. To account for this, some studies have normalized power in each frequency band to total spectral power to measure relative power^13,14,26^. While this approach reveals elevated relative delta power in AS, it introduces its own interpretive challenges: the dominance of low-frequency power skews the distribution of relative power, artificially suppressing higher-frequency bands and obscuring potentially meaningful differences outside the delta range^15,29^. Moreover, neither raw nor relative power metrics differentiate between genuine oscillatory (periodic) activity and broadband changes in the underlying 1/f-like (aperiodic) structure of the EEG signal, a biologically meaningful component in its own right^30^. Spectral parameterization^31^ (*specparam*) addresses these limitations by modeling power spectra as the sum of periodic and aperiodic components, enabling more accurate quantification of oscillatory power in specific frequency bands while separately characterizing the aperiodic structure. This relatively new approach has rapidly gained traction, with recent applications in tracking developmental changes in aperiodic and oscillatory activity^32^ and widespread use across studies of neurological and neurodevelopmental disorders^30^.

In this study, we applied spectral parameterization to EEG recordings from children with AS and LFP recordings from *Ube3a^m-/p+^*mice to investigate the electrophysiological basis of elevated delta power in AS. We separately analyzed periodic and aperiodic components of the power spectrum, then examined how these spectral features relate to behavioral phenotypes in both species and how they vary with age. Overall, our data dissect the underlying components of elevated delta power in AS, revealing distinct periodic and aperiodic signatures that map onto different behaviors and developmental windows. This work refines an established EEG biomarker and introduces aperiodic spectral features as a novel, translational marker for AS that may inform future clinical trials and preclinical research.

## Results

### Periodic and aperiodic spectral features contribute to elevated delta power in AS

We first quantified delta oscillations using standard spectral analysis methods using an existing dataset^29^ comprising 159 EEGs from children with AS (1-15 y) and 185 EEGs from age-matched typically-developing (TD) controls during periods of wake. As expected, we observed a distinct delta (2-4 Hz) peak in the power spectral densities (PSDs) of AS EEGs, which was quantified as an increase in raw delta power relative to TD controls (*t*(158.1) = 8.79, *p* < 0.001, *d* = 2.67; **Fig 1A-B**). As previously reported^15^, this increase in raw delta power was accompanied by a general elevation in broadband power (total power from 1-50 Hz: *t*(159.3) = 9.91, *p* < 0.001, *d* = 2.33), with significantly higher raw power also observed in the theta (4-6 Hz; *t*(159.2) = 11.54, *p* < 0.001, *d* = 2.17), alpha (6-12 Hz; *t*(312.7) = 6.62, *p* < 0.001, *d* = 0.81), beta (13-30 Hz; *t*(199.4) = 10.24, *p* < 0.001, *d* = 1.31), and gamma (30-50 Hz; *t*(244.2) = 6.42, *p* < 0.001, *d* = 0.95) bands (**Fig. 1B**). Analyzing relative power confirmed elevated relative delta and theta power in AS EEGs, but the dominance of low frequencies in the relative power calculation led to an artificial reduction in relative power in the alpha, beta, and gamma bands (**Fig S1**).

**Figure 1.**
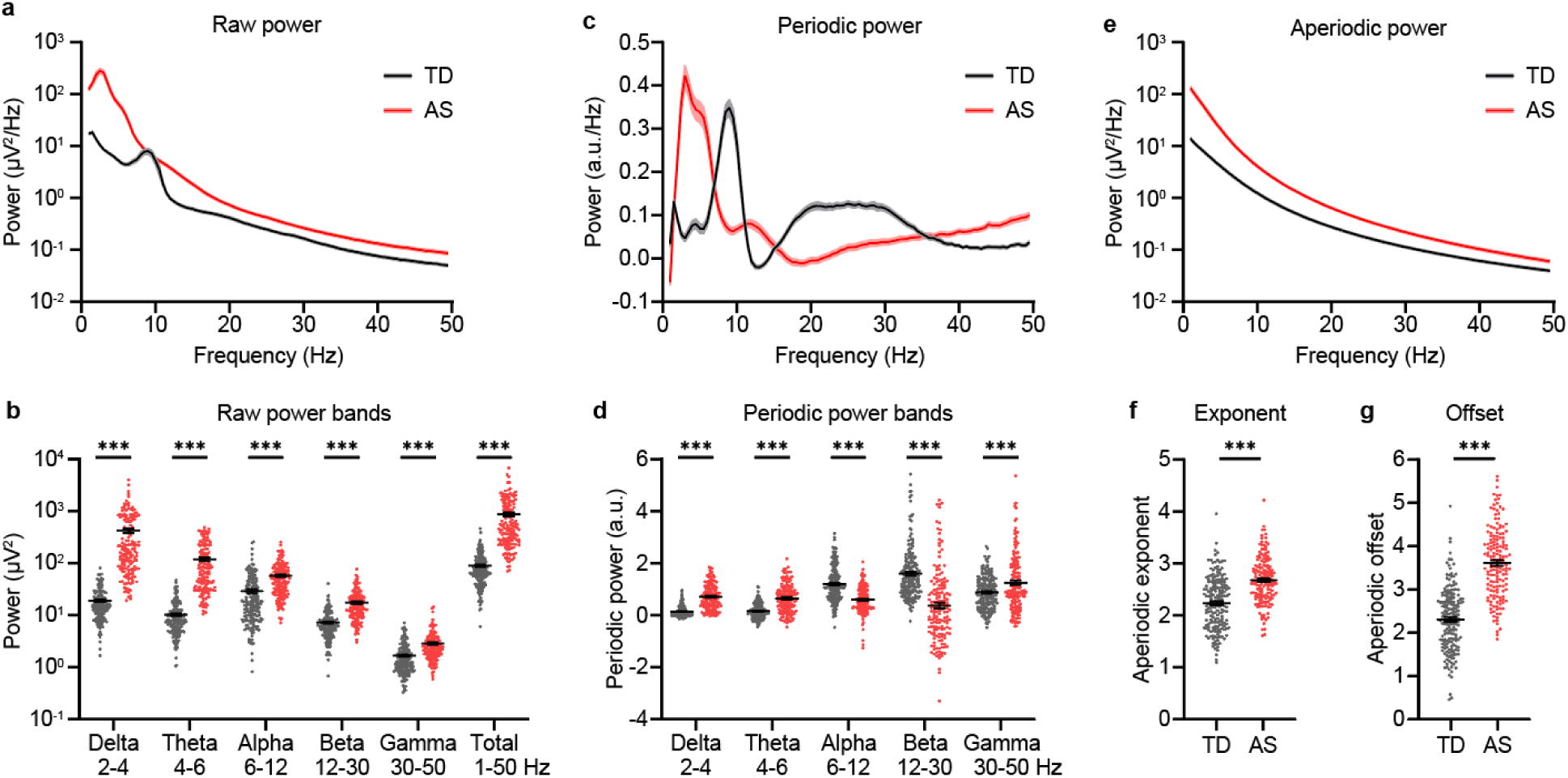
Periodic and aperiodic power spectra reveal group differences between TD and AS EEGs. **a.** Group PSDs from wake EEGs for all TD (black) and AS (red) EEGs. **b.** Average power is elevated across all frequency bands for AS EEGs. **c.** Group periodic power spectra calculated using *specparam*. **d.** Periodic delta, theta, and gamma power are increased in AS, and periodic alpha and beta power are decreased in AS. **e.** Group aperiodic spectra for TD and AS EEGs. Aperiodic exponent (**f**) and offset (**g**) are increased in AS EEGs. TD: *n* = 185 recordings/subjects, AS: *n* = 159 recordings from 95 subjects. **a,c,e:** Shading indicates ± SEM. **b,d,f,g:** Points represent individual recordings, black lines indicate group averages ± SEM. \*\*\**p* < 0.001, unpaired t-tests with Welch’s correction, followed by Holm-Šídák correction for multiple comparisons.

To overcome the limitations of raw and relative delta power measures, and to isolate changes in the underlying structure of the power spectra, we used spectral parameterization^31^ to separate EEG power spectra into periodic oscillatory peaks and an aperiodic 1/f-like background. For each EEG, we estimated the aperiodic component and subtracted it from the raw PSD to isolate periodic activity, revealing group differences in both the periodic and aperiodic power spectra. In the periodic spectra (**Fig 1C**), AS EEGs displayed a pronounced increase in periodic delta power compared to TD controls (*t*(209.6) = 14.08, *p* < 0.001, *d* = 1.61; **Fig 1D**). We also observed elevated periodic theta (*t*(231.7) = 10.35, *p* < 0.001, *d* = 0.56) and periodic gamma power (*t*(240.3) = 3.72, *p* < 0.001, *d* = 0.41), and reduced periodic alpha (*t*(337.3) = 9.30, *p* < 0.001, *d* = 0.75) and periodic beta power (*t*(278.8) = 9.14, *p* < 0.001, *d* = 1.11) in the AS cohort (**Fig. 1D**). The aperiodic component of the power spectra (**Fig 1E**) also differed substantially between groups. Relative to TD controls, AS aperiodic spectra had significantly steeper slopes (i.e. larger aperiodic exponent, *t*(341.3) = 8.61, *p* < 0.001, *d* = 0.95; **Fig 1F**) and higher offsets (*t*(320.6) = 15.56, *p* < 0.001, *d* = 1.27; **Fig 1G**). These changes were not limited to wakefulness: in a subset of participants with sleep EEG recordings, periodic delta power, aperiodic exponent, and aperiodic offset were all similarly elevated in AS (**Fig S2**). Together, these findings demonstrate that the increase in raw delta power in AS EEGs reflects the contributions of both elevated periodic delta oscillations and systematic shifts in the aperiodic background of the EEG signal.

### Periodic delta and aperiodic exponent differentially predict cognitive and motor deficits in AS

Prior studies have shown that delta power is correlated with clinical severity in AS^17,18^, but it is unclear whether these associations are driven by changes in periodic delta oscillations, aperiodic broadband shifts, or both. To address this question, we assessed the relationships between raw delta power, periodic delta power, aperiodic exponent, and aperiodic offset with cognitive performance, as measured by the Bayley-III Cognitive Scale, in the subset of 47 AS participants for whom both EEG and behavioral data were available. Twenty of these subjects had multiple Bayley scores and EEG recordings across different time points (**Fig S3A**), resulting in 88 total Bayley-EEG pairs. As previously reported^17,33–35^, Bayley cognitive scores (BCS) in AS improved with age and were lower in children with a deletion phenotype (**Fig S3B**). To evaluate relationships between BCS and EEG measures while accounting for within-subject variability, age, and genotype, we employed the mixed effects modelling framework previously validated by Ostrowski et al.^17^:

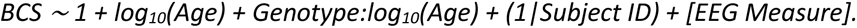

We evaluated separate models for each EEG measure (raw delta power, periodic delta power, aperiodic exponent, and aperiodic offset) and compared them to a partial model that excluded the EEG term. Models including raw delta power (**Fig 2A**) and periodic delta power (**Fig 2B**) significantly improved prediction of BCS relative to the partial model (partial model: AICc = 566.93; + raw delta power: ΔAICc = −15.45, *p* < 0.001; + periodic delta power: ΔAICc = −12.74, *p* < 0.001; **Table 1**). In contrast, models including the aperiodic exponent (**Fig 2C**) and offset (**Fig 2D**) did not yield significant improvements (+ aperiodic exponent: ΔAICc = −1.96, *p* = 0.11, + aperiodic offset: ΔAICc = −0.72, *p* = 0.17; **Table 1**). These results suggest that the predictive utility of raw delta power for cognitive deficits in AS is largely driven by its periodic component.

**Figure 2.**
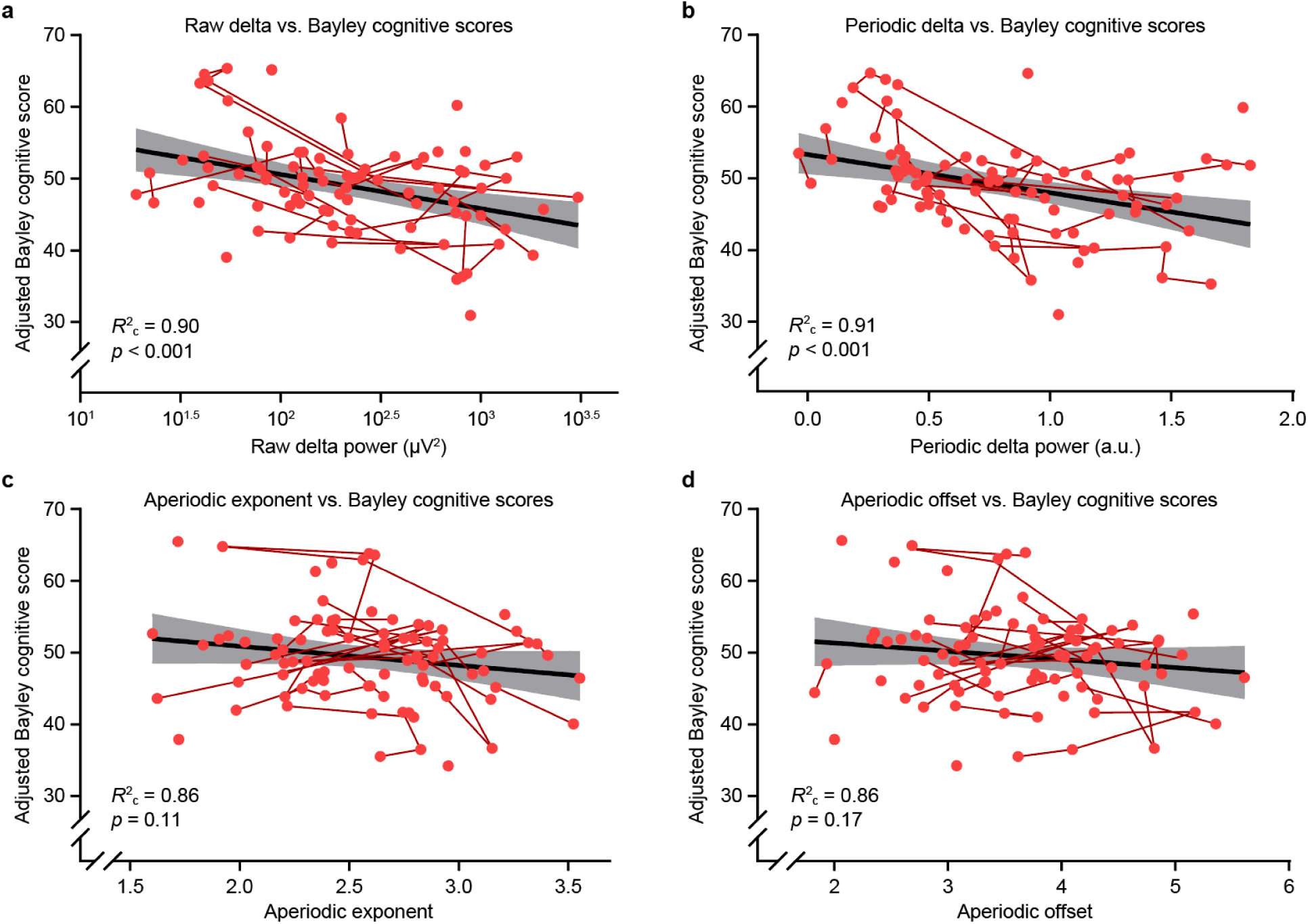
Raw and periodic delta power are predictive of cognitive ability in AS. Linear mixed-effects models (*BCS* ∼ 1 + log_10_ log_10_ *age* : *genotype* + (1|*ID*) + [*EEG measure*]) were used to relate Bayley cognitive scores (BCS) to each EEG measure, controlling for age (log_10_ *age*), age-genotype interactions (log_10_ *age* : *genotype*), and repeated measures (1|*ID*). **a-d.** Estimated Bayley cognitive scores, derived from the full model for each EEG measure, are plotted against raw delta power (**a**), periodic delta power (**b**), aperiodic exponent (**c**), and aperiodic offset (**d**). Each point represents a single recording; red lines connect repeated measures from the same subject. Black lines show linear fits with shaded 95% confidence intervals. Conditional *R*^2^ (*R*^2^c) values indicate fit of the full mixed model. *p* values are from likelihood ratio tests comparing full models to reduced models excluding the EEG measure. Bayley Cognitive Scores were adjusted to remove the modeled effects of age and genotype, allowing visualization of their relationship with delta power independent of these covariates.

**Table 1.** Comparisons of mixed-effects models predicting Bayley subdomain scores from EEG spectral features in children with Angelman syndrome. Each full model included one z-scored EEG predictor (Raw Delta, Periodic Delta, Aperiodic Exponent, or Aperiodic Offset), along with covariates log10(age), log10(age) × genotype, and a random intercept for subject ID. The first row in each column shows the corrected Akaike Information Criterion (AICc) for the partial model excluding EEG predictors. For each EEG feature, ΔAICc indicates the change in model fit relative to the partial model. Standardized regression coefficients (β ± SE) represent the change in raw Bayley subdomain scores per 1 standard deviation increase in the EEG feature. *p*-values indicate whether each EEG feature’s regression coefficient differs from zero, adjusted for multiple comparisons using the Benjamini-Hochberg method. Green-shaded cells indicate EEG features that significantly improved model fit (adjusted *p* < 0.05).

| Model | Bayley Subdomain |  |  |  |  |
| --- | --- | --- | --- | --- | --- |
|  | Cognitive | Receptive Communication | Expressive Communication | Fine Motor | Gross Motor |
| Partial Model | AICc: 566.93 | AICc: 472.62 | AICc: 419.84 | AICc: 499.90 | AICc: 553.21 |
| + Raw Delta | $\Delta\text{AICc: -15.45}$<br>$\beta: -2.79 \pm 0.61$<br>$p < 0.001$ | $\Delta\text{AICc: -1.91}$<br>$\beta: -0.87 \pm 0.41$<br>$p = 0.11$ | $\Delta\text{AICc: -0.84}$<br>$\beta: -0.55 \pm 0.31$<br>$p = 0.17$ | $\Delta\text{AICc: -3.26}$<br>$\beta: -1.15 \pm 0.48$<br>$p = 0.09$ | $\Delta\text{AICc: 2.13}$<br>$\beta: -0.29 \pm 0.65$<br>$p = 0.66$ |
| + Periodic Delta | $\Delta\text{AICc: -12.74}$<br>$\beta: -3.33 \pm 0.75$<br>$p < 0.001$ | $\Delta\text{AICc: 1.66}$<br>$\beta: -0.40 \pm 0.50$<br>$p = 0.53$ | $\Delta\text{AICc: 0.80}$<br>$\beta: -0.46 \pm 0.37$<br>$p = 0.31$ | $\Delta\text{AICc: 2.01}$<br>$\beta: -0.32 \pm 0.58$<br>$p = 0.65$ | $\Delta\text{AICc: 1.17}$<br>$\beta: -0.84 \pm 0.79$<br>$p = 0.38$ |
| + Aperiodic Exponent | $\Delta\text{AICc: -1.96}$<br>$\beta: -1.22 \pm 0.59$<br>$p = 0.11$ | $\Delta\text{AICc: -1.32}$<br>$\beta: -0.68 \pm 0.35$<br>$p = 0.14$ | $\Delta\text{AICc: 0.06}$<br>$\beta: -0.4 \pm 0.26$<br>$p = 0.22$ | $\Delta\text{AICc: -7.54}$<br>$\beta: -1.27 \pm 0.39$<br>$p = 0.012$ | $\Delta\text{AICc: 1.79}$<br>$\beta: -0.42 \pm 0.56$<br>$p = 0.54$ |
| + Aperiodic Offset | $\Delta\text{AICc: -0.72}$<br>$\beta: -1.08 \pm 0.61$<br>$p = 0.17$ | $\Delta\text{AICc: 0.02}$<br>$\beta: -0.56 \pm 0.37$<br>$p = 0.22$ | $\Delta\text{AICc: 0.72}$<br>$\beta: -0.35 \pm 0.27$<br>$p = 0.31$ | $\Delta\text{AICc: -2.48}$<br>$\beta: -0.93 \pm 0.42$<br>$p = 0.11$ | $\Delta\text{AICc: 2.07}$<br>$\beta: 0.28 \pm 0.58$<br>$p = 0.66$ |

To evaluate whether spectral features also predicted clinical severity in other behavioral domains, we applied the same modeling framework to receptive communication, expressive communication, fine motor, and gross motor scores from the Bayley-III. Most models did not show statistically significant improvements over the corresponding partial model after correcting for multiple comparisons (**Table 1**). The exception was the model including aperiodic exponent as a predictor of fine motor scores, which demonstrated a modest but statistically meaningful improvement over the corresponding partial model (partial model: AICc = 499.90; + aperiodic exponent: ΔAICc = −7.54, *p* = 0.012), indicating that increased aperiodic exponent was associated with decreased fine motor performance. Notably, periodic delta power was not predictive of fine motor scores (+ periodic delta power: ΔAICc = 2.01, *p* = 0.65). None of the EEG features significantly improved model fit for receptive communication, expressive communication, or gross motor outcomes.

### Developmental trajectories and regional assessment of spectral features in AS

We next examined how raw delta power, periodic delta power, aperiodic exponent, and aperiodic offset change across development in children with AS and TD controls. Using generalized additive mixed models, we modeled nonlinear trajectories of raw delta power, periodic delta power, aperiodic exponent, and aperiodic offset as a function of age for both AS and TD subjects (**Fig 3**). As expected^14,15^, we confirmed that raw delta power in children with AS declines with age, while remaining low and stable in TD controls (**Fig 3A**). However, the relatively steady decrease in raw delta power with age in AS appeared to reflect distinct developmental changes in periodic delta oscillations and in the aperiodic component of the EEG signal. Periodic delta power declined sharply in both groups during early childhood (∼ages 1 to 4) before stabilizing, but remained significantly elevated in AS into adolescence (**Fig 3B**). In contrast, aperiodic exponent and offset started low and increased from around 1 to 6 years old in TD children before steadying, whereas in AS children these parameters remained flat and elevated relative to TD controls across development (**Fig 3C-D**). These persistent group differences in both periodic and aperiodic features likely contribute to the sustained elevation of raw delta power in AS across the age range studied here.

**Figure 3.**
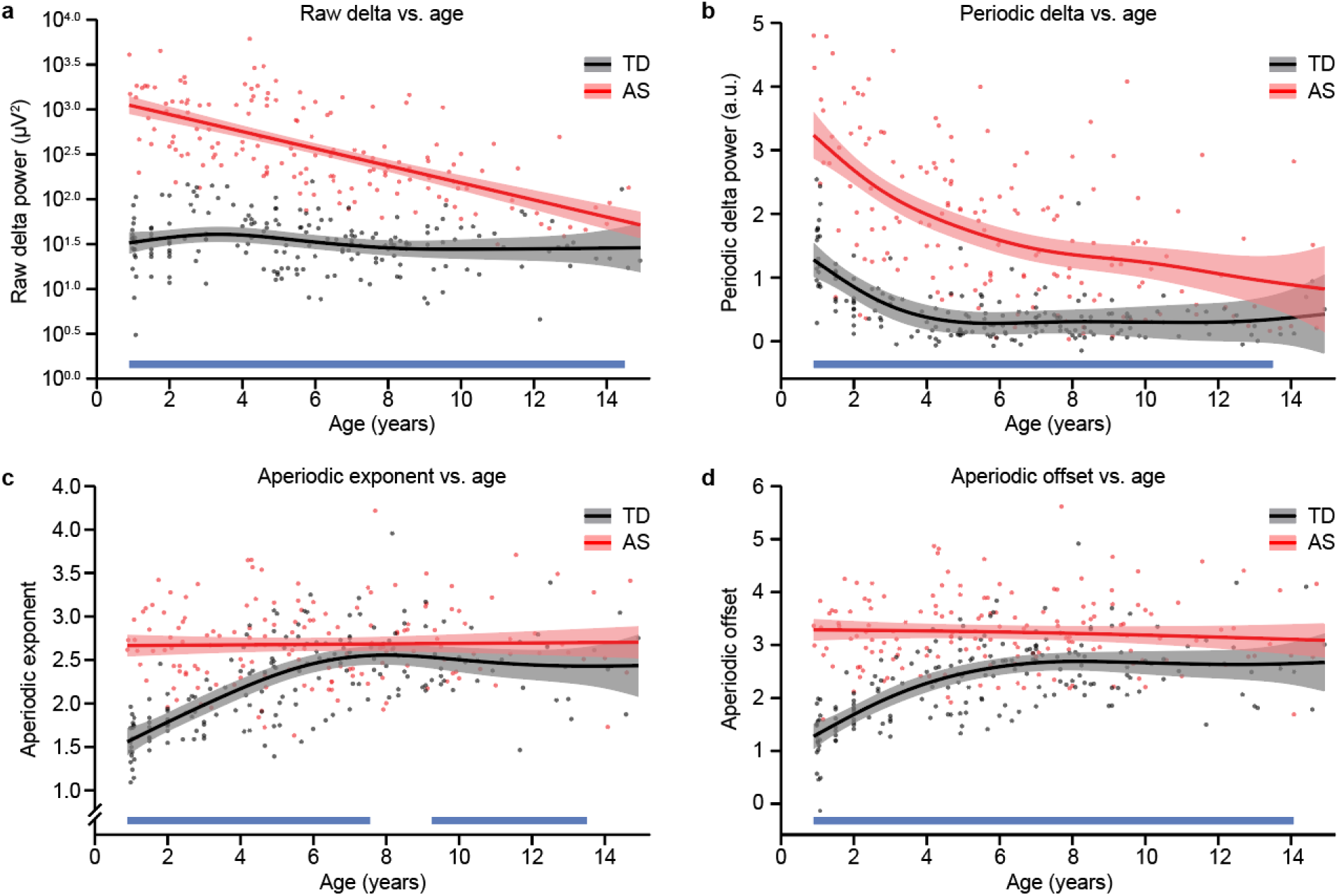
Developmental trajectories of periodic and aperiodic EEG features in AS. Generalized additive mixed model (GAMM) trajectories of raw delta power (**a**), periodic delta power (**b**), aperiodic exponent (**c**), and aperiodic offset (**d**) for TD (black) and AS (red) EEGs. Individual points correspond to single recording sessions. Lines represent predicted values for each group; shaded areas indicate 95% confidence intervals. Blue bars along the x-axis denote age intervals with statistically significant group differences (*p* < 0.05).

To examine regional variation in AS EEG phenotypes, we next compared raw delta power, periodic delta power, aperiodic exponent, and aperiodic offset across frontal, central, and occipital regions in AS and TD EEGs using linear mixed-effects models. All four measures showed significant group differences within each region (**Fig S4**). To assess regional variation in effect size, we calculated region-specific Cohen’s *d* values and compared these across regions. Effect sizes were similar across brain areas for raw delta power, trended larger frontally and centrally for periodic delta power, were largest occipitally for aperiodic exponent, and showed no regional variation for aperiodic offset (**Table S1**). These results suggest that although elevated delta power is broadly distributed, some of its spectral components, particularly aperiodic exponent, may show subtle regional variation.

### LFP recordings in AS model mice reveal conserved periodic and aperiodic spectral alterations

To evaluate whether the spectral alterations observed in children with AS are recapitulated in a preclinical mouse model, we analyzed LFP recordings from the primary visual cortex (V1) of *Ube3a^m-/p+^* mice and wild-type (WT, *Ube3a^m+/p+^*) littermate controls. Adult (P90) *Ube3a^m-/p+^* mice exhibited increased LFP power across a broad range of frequencies compared to WT controls (**Fig 4A**), with significant increases in raw power in the delta (3-5 Hz: *t*(25.17) = 5.08, *p* < 0.001, *d* = 1.47), theta (5-10 Hz: *t*(26.05) = 4.14, *p* = 0.001, *d* = 1.20), and beta (13-30 Hz: *t*(33.53) = 3.21, *p* = 0.006, *d* = 0.94) frequency ranges, while raw gamma power (30-50 Hz: *t*(36.33) = 0.41, *p* = 0.69, *d* = 0.12) did not differ by genotype (**Fig 4B**). To determine whether these power increases reflected changes in periodic oscillatory activity, background aperiodic activity, or both, we applied spectral parameterization using the *specparam* algorithm, similar to the human EEG analysis. Spectral parameterization revealed significant alterations in both periodic and aperiodic activity for adult *Ube3a^m-/p+^* mice relative to WT controls. In the periodic domain (**Fig 4C**), adult *Ube3a^m-/p+^* mice showed significantly elevated periodic delta (*t*(42.43) = 4.58, *p* = 0.001, *d* = 1.36) and beta (*t*(36.17) = 3.66, *p* = 0.002, *d* = 1.07) power, reduced periodic gamma power (*t*(42.32) = 10.34, *p* < 0.001, *d* = 3.09), with no significant change in periodic theta power (*t*(38.98) = 1.12, *p* = 0.27, *d* = 0.34, **Fig 4D**). Aperiodic spectra were steeper and shifted upward in adult *Ube3a^m-/p+^* mice (**Fig 4E**), reflected by increases in both the aperiodic exponent (*t*(36.85) = 7.59, *p* < 0.001, *d* = 2.23, **Fig 4F**) and offset (*t*(41.70) = 6.42, *p* < 0.001, *d* = 1.90, **Fig 4G**). Changes in raw delta, periodic delta, aperiodic exponent, and aperiodic offset were observed in both male and female adult *Ube3a^m-/p+^*mice (**Fig S5**).

**Figure 4.**
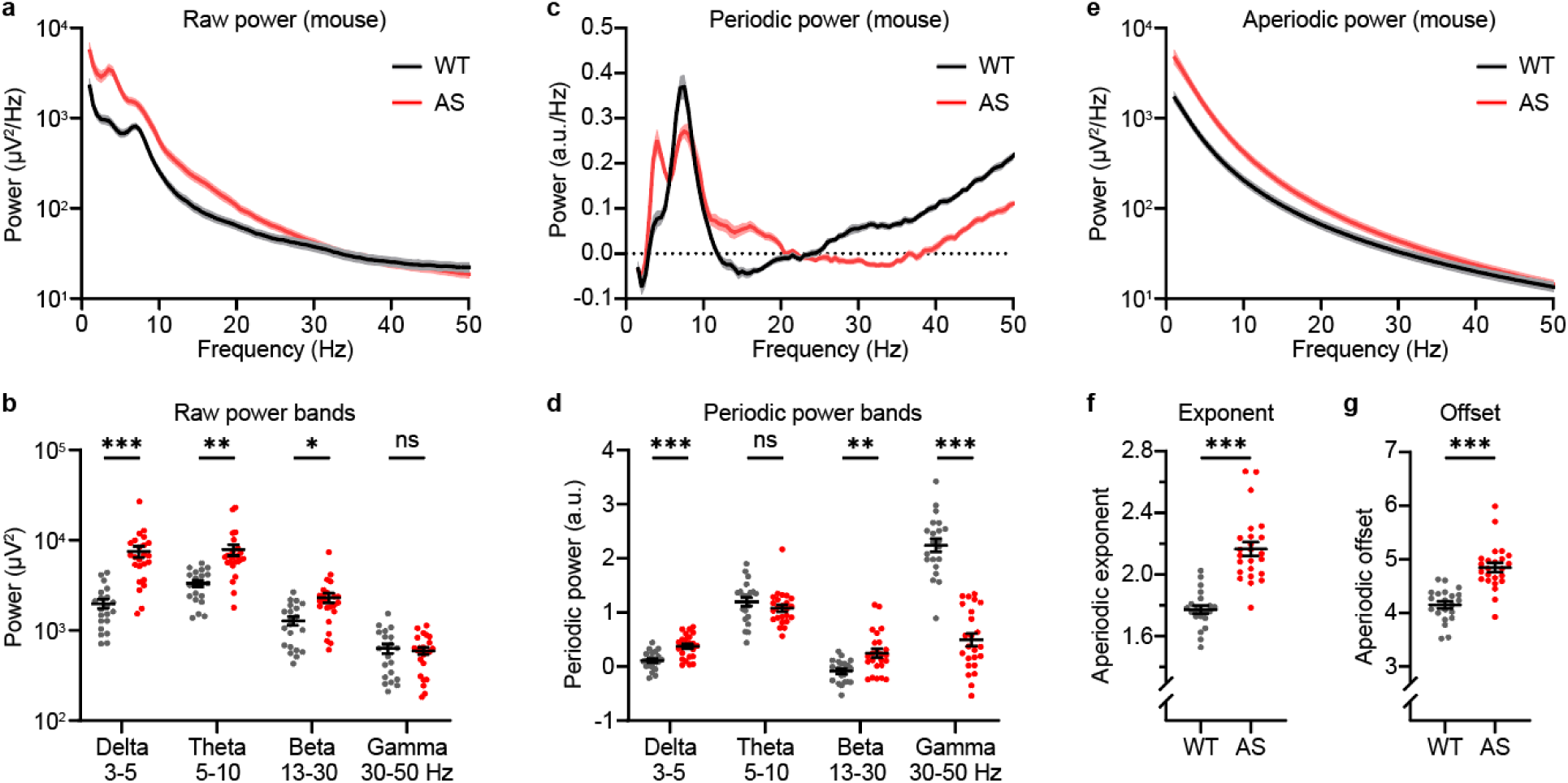
Spectral power alterations in V1 LFP recordings from Ube3a^m-/p+^ (AS) mice. **a.** Group-averaged power spectral densities (PSDs) from primary visual cortex (V1) recordings in WT (black, *n* = 21) and AS (red, *n* = 24) mice at P90. Shading indicates ± SEM. **b.** Average band-limited power is significantly increased in AS mice in the delta (3-5 Hz), theta (5-10 Hz), and beta (13-30 Hz) ranges, but not in the gamma band (30-50 Hz). **C.** Aperiodic power spectra extracted using spectral parameterization. **d-e.** Aperiodic exponent (**d**) and offset (**e**) are significantly elevated in AS mice. **f.** Periodic power spectra computed by subtracting the aperiodic fit from PSDs. **g.** AS mice show significantly elevated periodic delta and beta power, reduced gamma power, and no significant change in periodic theta power. In **b, d, e,** and **g**, each point represents an individual mouse; black lines indicate group averages. *p* values reflect unpaired t-tests with Welch’s correction, followed by Holm-Šídák correction for multiple comparisons (for **b** and **d**): * *p* < 0.05, ** *p* < 0.01, *** *p* < 0.001.

### Aperiodic spectral features are associated with behavioral deficits in Ube3a^m-/p+^ mice

To determine whether the spectral alterations observed in *Ube3a^m-/p+^* mice relate to behavioral impairments, we next examined correlations between LFP features and performance on an established behavioral battery for assessing *Ube3a^m-/p+^* phenotypes^36,37^. Prior to the LFP recordings at P90, mice underwent five days of rotarod testing, followed by open field, marble burying, and nest building assays (**Fig 5A**). As expected, *Ube3a^m-/p+^*mice showed behavioral impairments for all measures except open field center time (rotarod day 1: *t*(49.83) = 3.48, *p* = 0.001; rotarod day 5: *t*(40.23) = 3.49, *p* = 0.001; open field total distance: *t*(50.57) = 4.33, *p* < 0.001; open field center time: *t*(36.84) = 0.84, *p* = 0.4078; marble burying: *t*(50.78) = 11.25, *p* < 0.001; nest building: *t*(32.53) = 5.80, *p* = 0.001; **Fig 5B-G**). To summarize behavioral performance across multiple measures, we used principal component analysis (PCA) following the approach described by Tanas et al.^37^, accounting for sex differences in a subset of behaviors (**Fig. S6**). The first principal component (PC1) captured the majority (86.6%) of the genotype-related variance across all components (**Fig 5H, Table S1**) and was therefore used as a composite index of behavioral phenotype severity.

**Figure 5.**
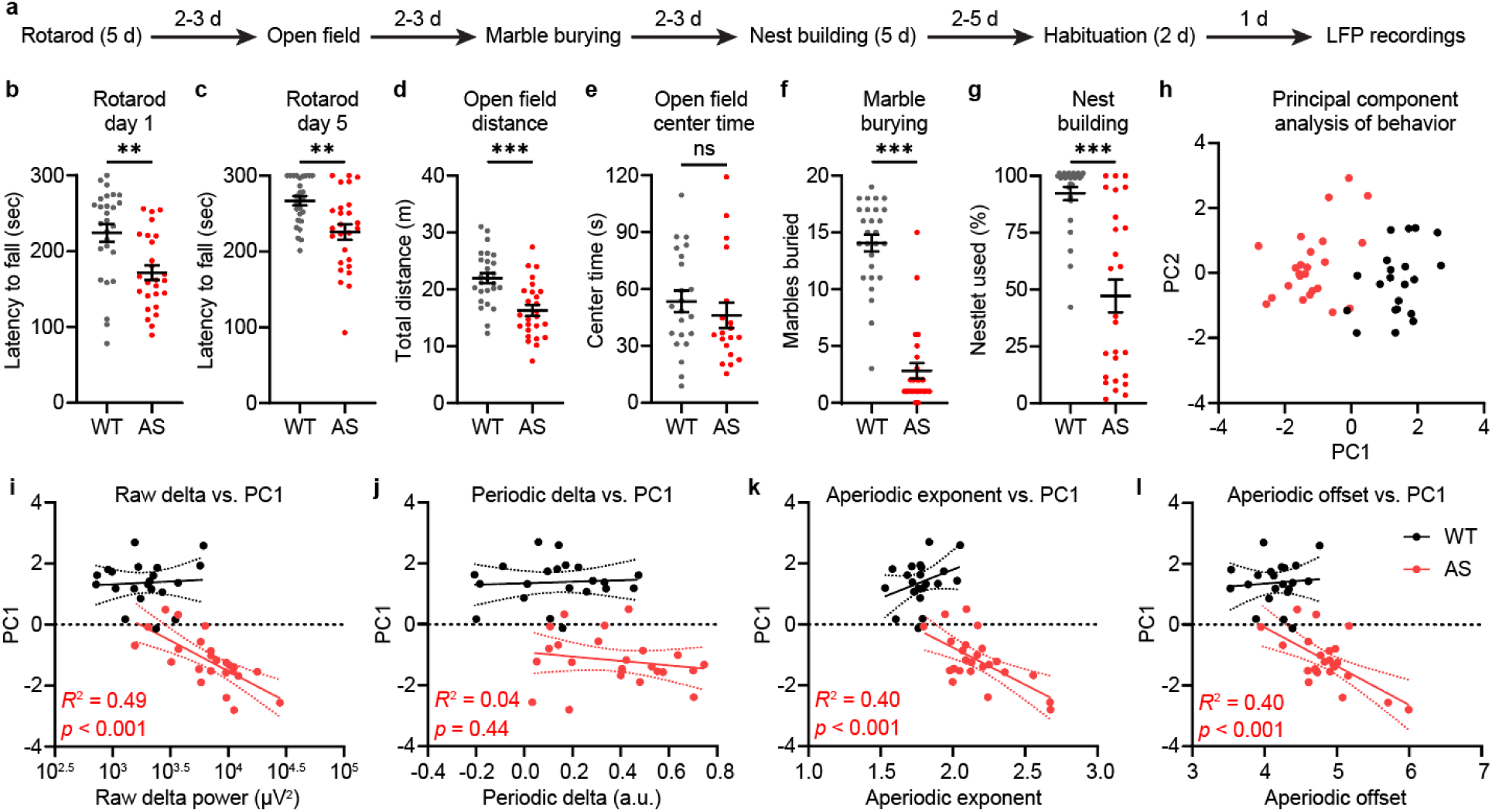
Aperiodic spectral features predict motor-related deficits in Ube3a^m-/p+^ (AS) mice. **a.** Timeline of behavioral testing preceding the final LFP recording. One cohort of mice was implanted at P54, preceding the behavioral battery, and a second cohort was implanted at P84, after the battery (see **Methods**). Both groups were recorded at P90 and data was pooled between them. **b-g.** Behavioral performance in WT (gray) and AS (red) mice across behavioral tasks. AS mice showed reduced performance on rotarod days 1 (**b**) and 5 (**c**), traveled shorter distances in the open field (**d**), buried fewer marbles (**f**) and used less nesting material (**g**). No difference was observed in open field center time (**e**). Black lines show mean ± SEM for each group. Asterisks show results from unpaired t-tests with Welch’s correction: \*\**p* < 0.01, \*\*\**p* < 0.001. **h.** Principal component analysis (PCA) of all six behavioral measures. PC1 explained the largest proportion of variance and separated WT from AS mice. **i-l.** In AS mice, PC1 was significantly associated with raw delta power (**i**), aperiodic exponent (**k**), and aperiodic offset (**l**), but not periodic delta power (**j**). Solid lines show linear fits; dashed lines indicate 95% confidence intervals. Reported *p* and R^2^ values show linear regression results for AS mice; no significant relationships were observed in WT mice (WT raw delta: *R*^2^ = 0.005, *p* = 0.766; periodic delta: *R*^2^ = 0.005, *p* = 0.772; aperiodic exponent: *R*^2^ = 0.114, *p* = 0.135; aperiodic offset: *R*^2^ = 0.007, *p* = 0.716).

Raw delta power was significantly correlated with PC1 in *Ube3a^m-/p+^*mice (*F*(1,22) = 19.73, *p* < 0.001, *R*^2^ = 0.47, **Fig 5I**), such that higher delta power was associated with greater behavioral impairment. To identify the source of this association, we separately examined correlations between PC1 and periodic delta power, aperiodic exponent, and aperiodic offset. This analysis revealed that the association was primarily driven by aperiodic features: periodic delta power was not significantly correlated with PC1 (*F*(1,22) = 0.61, *p* = 0.44, *R*^2^ = 0.03, **Fig. 5J**), whereas both aperiodic exponent (*F*(1,22) = 12.88, *p* = 0.002, *R*^2^ = 0.37, **Fig 5K**) and offset (*F*(1,22) = 10.18, *p* = 0.004, *R*^2^ = 0.32, **Fig 5L**) were. Notably, none of the LFP measures were significantly correlated with PC1 in WT mice, indicating that these relationships were specific to AS model mice. Together, these findings suggest that behavioral deficits in motor-related tasks in *Ube3a^m-/p+^* mice are closely linked to the aperiodic structure of the LFP power spectra.

### Aperiodic phenotypes precede periodic delta abnormalities in Ube3a^m-/p+^ mice

To assess how periodic and aperiodic phenotypes change over development, we analyzed LFPs from multiple mouse cohorts recorded at postnatal days (P) 28, 42, 60, and 90. At P28, raw and periodic delta power, aperiodic exponent, and aperiodic offset did not significantly differ between genotypes (**Fig 6**). Genotype differences emerged at later ages, with the periodic and aperiodic components following distinct temporal trajectories. Raw delta power (main effect of genotype: *F*(1,85) = 22.05, *p* < 0.001; main effect of age: *F*(3,90) = 11.68, *p* < 0.001; genotype × age interaction: *F*(3,90) = 2.234, *p* = 0.090) was significantly higher in *Ube3a^m-/p+^* mice relative to WT controls at both P60 (Šídák’s multiple comparisons test, *p* < 0.001) and P90 (*p* < 0.001; **Fig 6A**). Periodic delta power (genotype: *F*(1,85) = 10.08, *p* = 0.002; age: *F*(3,90) = 0.85, *p* = 0.47; genotype × age: *F*(3,90) = 4.29, *p* = 0.007) showed a significant genotype difference only at P90 (*p* < 0.001; **Fig 6B**). In contrast, aperiodic exponent and offset diverged earlier. Aperiodic exponent (genotype: *F*(1,85) = 52.09, *p* < 0.001; age: *F*(3,90) = 20.84, *p* < 0.001; genotype × age: *F*(3,90) = 7.00, *p* < 0.001) was significantly elevated in *Ube3a^m-/p+^* relative to WT mice beginning at P42 (*p* < 0.001), and remained elevated at P60 (*p* < 0.001) and P90 (*p* < 0.001; **Fig 6C**). Aperiodic offset (genotype: *F*(1,85) = 32.82, *p* < 0.001; age: *F*(3,90) = 24.79, *p* < 0.001; genotype × age: *F*(3,90) = 5.75, *p* = 0.001) was similarly elevated starting at P42 (*p* = 0.004), with significant group differences persisting at P60 (*p* < 0.001) and P90 (*p* < 0.001; **Fig 6D**). These findings indicate that changes in the aperiodic component of the LFP emerge earlier in development than periodic delta abnormalities in *Ube3a^m-/p+^*mice.

**Figure 6.**
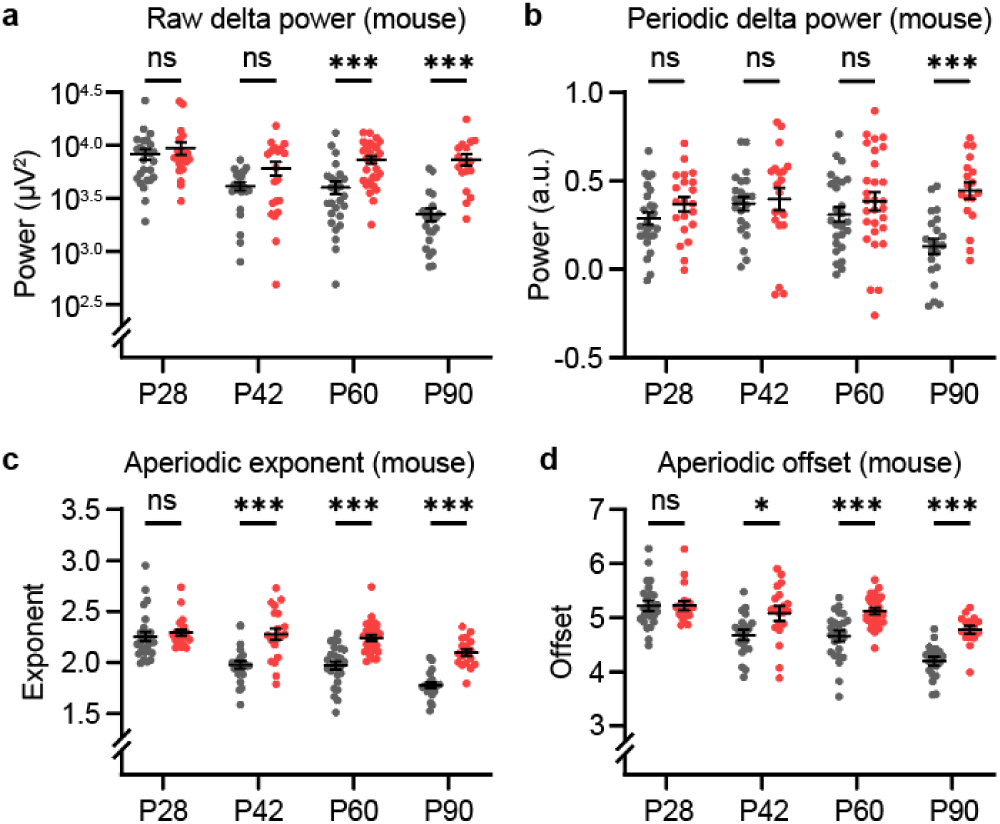
Developmental emergence of spectral alterations in Ube3a^m-/p+^ (AS) mice. Group comparisons of spectral features in WT (gray) and AS (red) mice across four developmental timepoints: P28, P42, P60, and P90. P90 data were previously shown in Fig 4 and are re-plotted here for comparison. **a.** Raw delta power shows no group difference at early ages but becomes significantly elevated in AS mice at P60 and P90. **b.** Periodic delta power is significantly increased in AS mice only at P90. **c-d.** Aperiodic exponent (**c**) and offset (**d**) are significantly higher in AS mice from P42 onward. *p*-values from Šídák’s multiple comparisons test. \**p*<0.05, \*\*\**p* < 0.001.

## Discussion

This study identified distinct periodic and aperiodic contributions to the elevated delta power phenotype characteristic of Angelman syndrome. Using spectral parameterization, we separated the rhythmic (periodic) and broadband (aperiodic) components of EEG recordings from children with AS and LFP recordings from *Ube3a^m-/p+^* mice. We found that increased delta power in both species reflects a combination of stronger periodic delta oscillations and a steepening and upward shift of the aperiodic background (**Figs 1, 4**). These spectral components were differentially associated with behavioral deficits (**Figs 2, 5**, **Table 1**) and diverged in their developmental trajectories (**Figs 3, 6**) for both mice and humans. Together, these findings establish periodic delta power and aperiodic spectral features as distinct and complementary biomarkers with high translational relevance.

While delta abnormalities have long been recognized in AS^11,38^, prior studies have typically quantified these changes using raw or relative power measures that do not distinguish between periodic oscillations and broadband spectral shifts^13,14,16–18,26^. Although these traditional approaches can reliably detect group-level differences (**Fig 1A-B, Fig S1**), they conflate separable spectral components with likely distinct physiological origins. By isolating the periodic component of the EEG signal, we more accurately quantified delta oscillations and found that both raw and periodic delta power were significantly associated with cognitive ability in AS (**Fig 2**, **Table 1**). Although raw and periodic delta power yielded similar improvements in model fit for predicting Bayley cognitive scores, periodic delta power likely reflects a more specific and physiologically interpretable marker of neural dysfunction in AS, as it excludes confounding contributions from broadband spectral activity.

Beyond revealing alterations in periodic delta power, our approach also quantified a previously uncharacterized aspect of the AS EEG phenotype: a broadband shift in the aperiodic component of the power spectrum, reflected by increases in aperiodic exponent and offset (**Fig 1E-G**). Although aperiodic exponent and offset are mathematically distinct parameters, they are often highly correlated due to the way they are estimated^39^. In our human dataset, only the aperiodic exponent was associated with fine motor deficits (**Table 1**), suggesting that it may be a more specific and behaviorally relevant biomarker than aperiodic offset for AS. This association aligns with growing evidence that the aperiodic exponent, or spectral slope, reflects clinically meaningful changes in neural function across a range of neurological and psychiatric conditions^30^. In addition, our finding that aperiodic exponent and periodic delta are linked to impairments in different behavioral domains and follow distinct developmental trajectories (**Fig 3**) indicates that these features likely reflect separable physiological mechanisms.

Prior studies using raw delta power have reported significant associations across multiple Bayley subdomains in children with AS^17,18^. We similarly found a robust relationship with cognitive scores, while other domains showed non-significant but directionally consistent effects. These differences likely reflect limited statistical power due to the need to correct for multiple comparisons across EEG features and behavioral domains. Together with prior findings, our results strengthen the evidence for delta power as a clinically meaningful biomarker in AS and further suggest that distinct spectral components may underlie different aspects of developmental function.

Given that periodic delta and aperiodic exponent are linked to distinct behavioral domains, separating these spectral features offers a more precise approach for evaluating treatment efficacy in AS clinical trials. Traditional EEG metrics that conflate periodic and aperiodic features may obscure subtle, domain-specific improvements. Spectral parameterization addresses this limitation by disentangling the underlying signals and enabling clearer interpretation of treatment-related changes. Because this method is open-source, user-friendly, and applicable to existing datasets, it offers a practical solution for both retrospective and prospective analyses. Incorporating both periodic and aperiodic measures into clinical trial pipelines may improve sensitivity to therapeutic effects and support the development of more targeted outcome metrics.

While periodic delta oscillations are relatively specific to AS^12,13^ and may reflect unique aspects of the disorder’s underlying physiology, alterations in aperiodic features have been reported across a range of neurodevelopmental and psychiatric conditions^30^. The widespread presence of aperiodic changes across disorders suggests that they may reflect common disruptions in circuit-level processes. In this context, studying aperiodic features in AS may help clarify how neural circuits are disrupted in AS and how those disruptions relate to patterns observed in other neurodevelopmental disorders. But importantly, biomarker specificity is not a requirement for evaluating treatment response; instead, robustness, quantifiability, and relevance to clinically meaningful outcomes are more important^40^. Both periodic delta and aperiodic exponent meet these criteria, supporting their complementary utility as EEG biomarkers in AS clinical trials.

Our findings also highlight the utility of spectral parameterization in preclinical models of AS, where reliable and translatable biomarkers are critical for drug development and mechanistic studies. In *Ube3a^m-/p+^* mice, periodic and aperiodic spectral changes closely mirrored those observed in humans (**Fig 4**). Notably, aperiodic exponent and offset phenotypes showed larger effect sizes than periodic delta power, diverged earlier from WT mice during development (**Fig 6C-D**), and were more strongly associated with motor-related behavioral deficits (**Fig 5K-I**), supporting their potential use as more sensitive and functionally relevant biomarkers in mice. Although periodic delta power was also elevated in *Ube3a^m-/p+^* mice (**Fig 4C-D**), it did not correlate with performance on the primarily motor-based tasks used in our behavioral battery (**Fig 5J**). This dissociation suggests that periodic delta power and aperiodic features may reflect distinct neural processes linked to different behavioral domains, mirroring their respective associations with cognition and motor function in children with AS. Future behavioral studies using more cognitively demanding tasks in *Ube3a^m-/p+^* mice^41–45^ may help clarify whether periodic delta power can serve as a cross-species marker of cognitive function in AS.

Despite broad similarities in the spectral phenotypes of humans and mice, we observed several notable differences that present challenges for cross-species translation. In children, periodic and aperiodic abnormalities were present from the earliest ages recorded and remained elevated relative to typically developing controls through at least 13 years of age (**Fig 3**). In contrast, *Ube3a^m-/p+^*mice diverged from WT controls later in development, with aperiodic differences emerging around P42 and periodic delta differences not appearing until P90 (**Fig 6**). This discrepancy may suggest potential species-specific differences in brain maturation or circuit development. Another challenge lies in mapping frequency bands between species. The elevated 3-5 Hz periodic peak observed in *Ube3a^m-/p+^* mice overlaps with the traditional human delta band but has also been proposed as a rodent analog of human alpha oscillations^46–48^, though this interpretation is debated^49^. These low-frequency changes could reflect homologous pathophysiology despite differences in frequency range, or might arise from distinct mechanisms in mice and humans. Clarifying the relationship between these species-specific oscillations will be critical for validating periodic delta power as a translational biomarker.

Our results offer a starting point for investigating the neural mechanisms underlying periodic and aperiodic alterations in AS. Delta-range oscillations are thought to arise from dynamic interactions within corticothalamic circuits, including thalamocortical relay neurons expressing HCN and T-type calcium channels^50–52^, GABAergic interneurons in the thalamic reticular nucleus^53^, and intrinsically bursting pyramidal neurons in neocortical layer 5^54,55^. In *Ube3a^m-/p+^*mice, increased low-frequency power depends specifically on Ube3a loss in GABAergic neurons^21^, suggesting that dysfunction within inhibitory elements of the corticothalamic circuit may drive elevated periodic delta power in AS. In contrast, the physiological basis of aperiodic spectral features remains less well understood. A steeper aperiodic slope (i.e. larger exponent) is often interpreted as reflecting a relative increase in inhibition^30,56,57^, but this interpretation conflicts with the cortical hyperexcitability observed in AS model mice^58–60^ and the high prevalence of seizures in individuals with AS^1^. These discrepancies suggest that aperiodic alterations may reflect more complex cortical network dysfunction not captured by simple excitation-inhibition models^61^. Future studies combining circuit-level manipulations with *in vivo* recordings will be useful for identifying the mechanisms underlying these spectral changes in AS.

Together, our findings offer a more nuanced understanding of the elevation in delta power observed in AS EEGs, with important implications for biomarker development. By disentangling periodic and aperiodic contributions to the delta phenotype, we clarify the electrophysiological features that comprise this useful biomarker and uncover distinct behavioral associations that may inform the design of clinical trials. The cross-species consistency of these features strengthens their translational value and supports their potential utility for evaluating treatment efficacy in clinical and preclinical settings. Future studies should aim to identify the cellular and circuit-level origins of these features, both to improve therapeutic targeting in AS and to deepen our understanding of how molecular and cellular alterations can disrupt network dynamics across neurodevelopmental disorders. Our findings suggest that future trials may benefit from evaluating periodic delta oscillations and aperiodic features as distinct, complementary biomarkers, each potentially capturing different aspects of neural dysfunction in AS.

## Methods

### Participants

We performed a retrospective analysis using a previously reported dataset^29^ containing deidentified EEG recordings from 95 children with AS (159 EEGs) and 185 age-matched typically developing (TD) children (one EEG each). Recording ages ranged from 0.9 to 14.9 years (AS = 6.1 ± 0.3 y; TD = 5.8 ± 0.3 y, mean ± SEM). The AS group included 38 females and 57 males; the TD group included 86 females and 99 males. Of the AS recordings, the majority (149 recordings from 88 unique subjects) came from the Angelman Natural History Study (ClinicalTrials.gov identifier: NCT00296764) conducted at Massachusetts General Hospital and shared via the LADDER database^62^. An additional 10 recordings from 7 subjects came from the UNC Sleep Disorders Center^13^. 35 AS participants (33 MGH, 2 UNC) contributed longitudinal data. Genotype information was available for 89 of the 95 AS children (67% deletion; 10% uniparental disomy; 4% imprinting defect; 15% UBE3A mutation; 3% abnormal methylation). TD recordings were collected at MGH (98), UNC (10) and UCLA (77). Children were classified as typically developing on the basis of normal developmental history and the absence of epilepsy or other neurological disorders.

### Neurodevelopmental testing

48 AS participants from the Natural History Study completed the Bayley Scales of Infant and Toddler Development, Third Edition, within two months of EEG acquisition (median interval = 0 days, range = 0-39 days). After removal of one child with incomplete results, the final dataset comprised 88 EEG-behavior pairs from 47 children; 20 contributed data at multiple visits. Raw domain scores (Cognitive, Expressive Communication, Receptive Communication, Fine Motor, Gross Motor) were used for analysis.

### EEG data collection and processing

EEG data in the present study were previously cleaned and processed for a separate study^29^. EEGs were collected during awake, task-free recordings for all subjects, and during sleep for a subset of subjects (**Fig S2**). Periods of sleep and wake were identified and annotated by a trained neurologist. Raw EEGs were preprocessed similar to as described in prior work^13,14^ using custom MATLAB scripts. Preprocessing steps included re-referencing to the common average, applying a 0.1 Hz high-pass filter, a 100 Hz low-pass filter, and a 60 Hz notch filter to remove noise and electrical interference. Recordings were then visually inspected to identify and exclude artifacts and excessively noisy channels. To ensure consistency between datasets, we analyzed six channels from three regions of interest: F3 and F4 (frontal), C3 and C4 (central), and O1 and O2 (occipital).

### EEG power spectral analysis

Power spectral densities (PSDs) for each channel of interest were computed in MATLAB (R2024a) with Welch’s method using 2-s Hamming windows and 50 % overlap, yielding 0.5 Hz resolution. Median PSDs for each electrode were converted to linear µV² units; raw power was summed within delta (2-4 Hz), theta (4-6 Hz), alpha (6-12 Hz), beta (13-30 Hz) and gamma (30-50 Hz) bands. These frequency ranges were selected to align with prior studies characterizing oscillatory activity across early development^32^. PSDs from each channel pair (F3/4, C3/4, and O1/2) were averaged to obtain EEG power for each region of interest, and averaged across all three regions to obtain the average PSD for each subject. Participants with data missing for both electrodes from any region were excluded from analysis (4 AS and 2 TD subjects excluded).

Periodic and aperiodic components of each PSD were separated using *specparam* v2.0.0-rc2 (formerly *FOOOF*)^31^ in Python 3.12. The *specparam* algorithm models the log-transformed spectrum as the sum of a broadband aperiodic background and a set of Gaussian peaks that capture periodic oscillations. The aperiodic component is fit to:

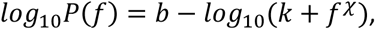

where *b* is the offset, 𝑘 is the knee parameter, and 𝜒 is the exponent (slope). The algorithm iteratively fits this background, subtracts it, identifies peaks on the residual, and refines both sets of parameters. Fits were performed from 1 to 49.5 Hz using the “knee” parameter, as low-frequency curvature was prominent in pilot fits and in earlier 1/f-corrected analyses of the same dataset^29^. Peak-detection settings were peak_width_limits = 0.5-12 Hz and peak_threshold = 2.0; no hard cap was placed on the number of peaks (max_n_peaks = inf), although empirical fits yielded only 3-10 peaks per recording. Model fit was excellent across subjects (average R^2^ ± *SD* for AS: 0.996 ± 0.005, TD: 0.997 ± 0.007). The knee frequency was calculated as 𝐹_𝑘𝑛𝑒𝑒_ = 𝑘^1/𝜒^, and was similar between groups (average Fknee ± *SD*, NT: 4.15 ± 1.92 Hz; AS: 3.22 ± 1.73 Hz). For each power spectrum, we extracted the aperiodic component using *specparam* and subtracted this component from the raw PSD to obtain the periodic component. Because the subtraction was performed in log space, the resulting periodic component reflects a power ratio rather than an absolute value, and is therefore unitless. Periodic power within each frequency band was calculated by summing within the same bands as above. Aperiodic exponent and aperiodic offset parameters were output from the *specparam* algorithm for each recording.

### Animals

All mouse protocols were approved by the Institutional Animal Care and Use Committee (IACUC) of Children’s National Medical Center. Mice were group-housed on a 12:12 light/dark cycle with ad libitum access to food and water. Male and female mice were used for experiments in equal genotypic ratios. Female *Ube3a^m+/p-^* × male *Ube3a^m+/p+^* breeders generated littermate experimental *Ube3a^m-/p+^* (AS) and *Ube3a^m+/p+^* wild-type (WT) mice. Mice were originally obtained from Jackson Labs (Bar Harbor, ME) (JAX #: 016590) and were maintained on a congenic C57BL/6 background.

### Surgeries and LFP recordings

Surgery and local field potential (LFP) recordings were conducted as previously described^14^ with minor alterations. Mice were anesthetized with 3% isoflurane. Before surgical incision, the head was shaved and the scalp cleaned with povidone-iodine (10% w/v) and ethanol (70% v/v). The scalp was resected and a steel head post was affixed to the skull (anterior to bregma) with cyanoacrylate glue. Small burr holes were drilled above both hemispheres of V1 (-3.6 mm AP, +/-3.0 mm ML relative to bregma). Tapered 300-500 kΩ tungsten recording electrodes (FHC), 75 µm in diameter at their widest point, were implanted in each hemisphere 470 µm below the cortical surface to target layer 4 of V1. Silver wire (A-M Systems) reference electrodes were implanted in the cerebellum. Electrodes were secured using cyanoacrylate, and the skull was covered with dental cement (Metabond). Nonsteroidal anti-inflammatory drugs were administered on return to the home cage (Carprofen, 5.0 mg/kg s.c.). Signs of infection and discomfort were carefully monitored. Mice were allowed to recover for at least 48h before head fixation and recording.

Mice were habituated to head-fixation over two days for 30 minutes each day, and then recorded for 20 minutes on each subsequent test day. During each recording session, mice viewed a static grey screen in an otherwise dark, quiet environment. Local field potentials (LFPs) were recorded using a Plexon OmniPlex Neural Recording Data Acquisition System. Signals were amplified and digitized at a 40 kHz acquisition rate, band-pass filtered from 0.1-200 Hz, and then down sampled to 1 kHz for analysis.

We originally aimed to record mice across all timepoints of interest (P28, P42, P60, and P90), but degradation of recording quality with age in some mice led to substantial attrition at the later time points. Therefore, we separated mice into 3 cohorts: cohort 1 (n=26 WT, 21 AS) was implanted at P22-23 and recorded at P28, P42, and P60; cohort 2 (n=13 WT, 12 AS) was implanted at P54-55 and recorded at P60 and P90; cohort 3 (n=8 WT, 6 AS) was implanted at P84-85 and recorded at P90. We did not observe significant differences between cohorts 1 and 2 at P60, or between cohorts 2 and 3 at P90, and therefore pooled all three cohorts together for our longitudinal analysis (see *Statistical analysis*).

### LFP analysis in mice

For each mouse, we analyzed data from the hemisphere with the better LFP signal (defined as having greater total power from 1-100 Hz). Recordings were binned into 10-second segments, and any segments with movement artifacts were manually removed. We analyzed PSD for all remaining segments with Welch’s method using the pwelch function in MATLAB (2 second window with 50% overlap) resulting in frequency bins of 0.5 Hz. The PSD was interpolated between 59 and 61 Hz to account for 60 Hz line noise, and the PSDs across all 10-second segments were averaged to get the average PSD for each mouse. Power within different frequency bands was calculated by integrating the PSD across the frequency range of interest. We defined delta as 3-5 Hz, theta as 5-10 Hz, beta as 13-30 Hz, and gamma as 30-50 Hz. We used this delta frequency range based on previous findings that *Ube3a^m-/p+^* mice on a C57BL/6 background show a stronger delta phenotype at this slightly higher frequency range^14^ and because this was the frequency range at which we saw a clear peak in the periodic power spectrum after applying spectral parameterization. The theta, beta, and gamma frequency ranges were chosen to be consistent with previous publications^14^.

PSDs were parameterized into periodic and aperiodic components using the MATLAB wrapper for the *specparam* Python package (https://github.com/fooof-tools/fooof_mat). Each PSD was fit with a knee from 1-100 Hz (max_n_peaks = 5, peak_width_limits = [1, 30], peak_threshold = 2.0) to obtain the aperiodic power spectrum. The aperiodic exponent (*χ*) and offset (*b*) were obtained from the aperiodic fitting function (see *EEG power spectral analysis*). The knee frequency was calculated as 𝐹_𝑘𝑛𝑒𝑒_ = 𝑘^1/𝜒^, and did not differ significantly between groups (average Fknee ± *SD*, WT: 3.23 ± 1.48 Hz; AS: 3.64 ± 1.64 Hz). The periodic power spectrum was calculated as log10(PSD) - log10(aperiodic power). Periodic power for each frequency band was calculated by integrating the periodic power spectrum across the frequency range of interest.

### Behavioral testing in mice

Mice were run through a standard series of behavioral tests for *Ube3a^m-/p+^* model mice^36,37^, done in the order of rotarod, open field, marble burying, and nest building. The behavior battery began at P61-P65 in cohorts 2 and 3, following LFP electrode implantation but prior to P90 LFP recordings, with 2-3 days in between each behavioral test (see **Fig 5** schematic). The forced swim test was excluded from the standard behavioral battery because the implanted electrodes impeded swimming ability and therefore introduced excessive risk.

*Rotarod.* Mice were placed on a rotating rod (Ugo-Basile model #47600) programmed to accelerate from 4 to 40 rpm over a 5-minute period at a rate of 7.2 rpm². Each trial began immediately after the mouse was placed on the rod. Trials ended when the mouse either fell off the rod, completed three full consecutive passive rotations, or reached the 300-second time limit. Mice completed two trials per day, separated by approximately four hours, over five consecutive days. The average latency to fall for the two daily trials was used as the outcome measure for each day.

*Open field*. Mice were placed individually in a 42 × 42 cm open field arena (AccuScan Instruments, Inc., Columbus, OH), and their activity was recorded for 20 minutes using an infrared beam grid system (Omnitech Electronics, Inc. SuperFlex Open Field System). Total distance traveled in the x-y plane was calculated based on interruptions of the infrared beams as the mice moved throughout the arena, and center time was calculated as the percentage of the time that mice spent in the center 26 × 26 cm square of the arena.

*Marble burying*. The marble burying test was conducted in individual 16 × 8 inch cages filled with ∼4 inches of corncob bedding (Bed-o’Cobs ¼” bedding). Twenty glass marbles were evenly spaced on the surface of the bedding in a 5 × 4 grid. Each mouse was placed in a separate cage for a 30-minute trial, after which the number of marbles buried was recorded. A marble was considered buried if at least 50% of its surface was covered by bedding at the end of the trial.

*Nest building*. Following marble burying, mice were weighed, singly housed, and habituated to the nesting material (Bio-Rad #1703965) for three days. After habituation, fresh pre-weighed nesting material was introduced on day 1 of a 5-day testing period. On day 5, the largest remaining piece of nesting material was weighed. If no solid piece weighing more than 0.1 g remained the remaining weight was scored as 0. Percent of nestlet used was calculated as the difference between the initial and final weights divided by the starting nestlet weight.

### Principal component analysis of behavior

Principal component analysis (PCA) of the mouse behavioral battery was carried out as described in Tanas et al.^37^ using the PUMBAA MATLAB package (https://github.com/sidorovlab/PUMBAA). Six behavioral measures were included: rotarod day 1, rotarod day 5, open field total distance, open field center time, marbles buried, and percent of nestlet used. Mouse weight was excluded to focus the analysis on behavioral and motor performance; however, an exploratory analysis including weight did not qualitatively affect the results (data not shown). To account for sex-related differences in behavior, behavioral measures that showed a significant main effect of sex or a sex × genotype interaction (rotarod days 1 and 5) were standardized by *z*-scoring separately for males and females^37^. Measures without significant sex effects (open field distance, open field center time, marbles buried, and percent nestlet used) were *z*-scored across all mice. PCA was then performed on the six standardized measures. We computed the variance explained by each principal component and examined their loading distributions (**Table S1**). To assess the extent to which the resulting principal components captured genotype-related variance, we fit linear models predicting each PC from genotype and computed the proportion of variance explained (R^2^, **Table S1**). The first principal component (PC1) exhibited relatively uniform loadings across behavioral domains and accounted for the largest proportion of genotype-related variance. PC1 was therefore used as a composite index of behavioral severity in *Ube3a^m-/p+^* mice.

### Statistical analysis and modeling

*LMMs for behavioral prediction from EEG*. To determine whether EEG-derived spectral features predict behavioral outcomes in children with AS (**Fig 2**), we used linear mixed-effects models (LMMs) for each subscale of the Bayley-III: cognitive, receptive communication, expressive communication, fine motor, and gross motor scores. All models were fit using the lmer function in the lme4 package (R version 4.4.2), with model comparisons and performance metrics computed using the lmerTest package.

For each behavioral outcome, we first fit a reduced model that included fixed effects for log10-transformed age (log_age) and an interaction between age and genotype (log_age × genotype), along with a random intercept for subject ID to account for repeated measures. For example:

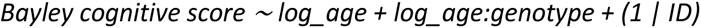

We then fit separate full models by adding one EEG measure (*z*-scored raw delta power, periodic delta power, aperiodic exponent, or aperiodic offset) as an additional fixed effect. For example, the full model predicting cognitive score from raw delta power was specified as:

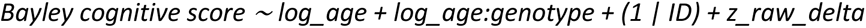

This modeling procedure was repeated separately for each Bayley subscale (cognitive, receptive communication, expressive communication, fine motor, and gross motor). Models were fit using maximum likelihood estimation, and evaluated using conditional R² values (R²c), along with the Akaike Information Criterion corrected for small sample sizes (AICc). Regression coefficients (β ± SE) and *p*-values were extracted for each EEG feature to quantify its predictive relationship with behavioral scores. *p*-values were computed from Wald t-tests using Satterthwaite’s approximation for degrees of freedom, and adjusted for multiple comparisons using the Benjamini-Hochberg false discovery rate (FDR) correction. To compare full models with their corresponding partial model, we calculated ΔAICc as AICcfull_model - AICcpartial_model, with ΔAICc ≤ -4 indicating a statistically meaningful improvement in fit over the partial model^63^.

*GAMMs for modeling age-related EEG changes*. To evaluate how EEG spectral features vary with age in children with AS compared to TD controls (**Fig 3**), we used generalized additive mixed models (GAMMs) to capture non-linear age effects while accounting for repeated measures within individuals. Models were implemented using the mgcv package in R (version 4.4.2). Each model was specified as:

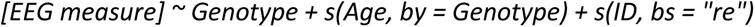

In this formula, Genotype indicates group membership (NT or AS), s(Age, by = Genotype) fits separate smooth functions of age for each genotype, and s(ID, bs = “re”) includes a random intercept for each participant to account for repeated measurements. Separate models were fit for each EEG measure: raw delta power, periodic delta power, aperiodic exponent, and aperiodic offset. Predicted trajectories were calculated by marginalizing over random effects and visualized across a continuous age range spanning the observed data. 95% confidence intervals were calculated from the standard errors of the model predictions. To identify time points with significant group differences, we computed pointwise *z*-statistics using the predicted values and their associated standard errors, and calculated two-tailed *p*-values assuming asymptotic normality. Age intervals with *p* < 0.05 were marked with blue bars beneath the plots.

*LMMs and effect size estimation for regional comparisons.* To assess genotype-related differences in spectral features across brain regions (**Table S1**, **Fig S4**), we used the lme4 package (R version 4.4.2) to fit separate LMMs for each EEG metric, including fixed effects for Genotype, Region, their interaction (Genotype × Region), and mean-centered Age, with a random intercept for Subject to account for repeated measures. Type-III tests were used to evaluate main and interaction effects.

For each region, we computed Cohen’s *d* to quantify standardized differences between NT and AS groups, using pooled standard deviations. Standard errors were calculated analytically, and estimated marginal means (emmeans package) were used to obtain genotype contrasts adjusted for covariates, assuming average age.

To test whether genotype effects varied by region, we computed pairwise differences in region-specific Cohen’s *d*, assuming independence. *Z*-tests were used to assess significance, and FDR correction (Benjamini-Hochberg) was used for multiple comparisons across region pairs.

*Other statistical comparisons*. For statistical comparisons of EEG and LFP measures (**Figs 1**, **4, S1, S2**) or behaviors (**Fig 5**) between groups, we used two-tailed unpaired *t*-tests with Welch’s correction. Multiple comparisons were corrected using the Holm-Šidák method. For analyses involving genotype and sex (**Figs S5, S6**), we used two-way ANOVAs with Tukey’s post hoc correction. For longitudinal recordings in mice involving multiple cohorts with different ages (**Fig 6**), we used a mixed effects model with genotype and age as fixed effects and animal ID as a random effect. The model was estimated using restricted maximum likelihood (REML), and Šídák’s multiple comparisons test was used for post hoc tests. For all analyses, uncorrected alpha was set to 0.05. Effect sizes are reported as Cohen’s *d* unless otherwise noted. Statistical analyses were performed with Prism 10 (GraphPad; RRID:SCR_002798).

## Acknowledgments

This work was supported by the Foundation for Angelman Syndrome Therapeutics (FT2002-003) (PI: MSS/AHD), the Angelman Syndrome Foundation (PI: MSS), NINDS T32NS115656 (DPM; PI: M. Berl), and by the District of Columbia Intellectual and Developmental Disabilities Research Center (DC-IDDRC) P50HD 105328 by NICHD (PI: W. Gaillard). We thank Dr. Catherine Chu and Katherine Walsh (MGH), Dr. Zheng Fan (UNC), and LADDER for providing data access, and Dr. Caleigh Guoynes (Children’s National) for assisting with open field behavioral testing.

## References

1 Thibert, R. L., Larson, A. M., Hsieh, D. T., Raby, A. R. & Thiele, E. A. Neurologic manifestations of Angelman syndrome. Pediatr Neurol 48, 271–279 (2013). 10.1016/j.pediatrneurol.2012.09.015

2 Tjeertes, J. et al. Enabling endpoint development for interventional clinical trials in individuals with Angelman syndrome: a prospective, longitudinal, observational clinical study (FREESIAS). J Neurodev Disord 15, 22 (2023). 10.1186/s11689-023-09494-w

3 Kishino, T., Lalande, M. & Wagstaff, J. UBE3A/E6-AP mutations cause Angelman syndrome. Nat Genet 15, 70–73 (1997). 10.1038/ng0197-70

4 Matsuura, T. et al. De novo truncating mutations in E6-AP ubiquitin-protein ligase gene (UBE3A) in Angelman syndrome. Nat Genet 15, 74–77 (1997). 10.1038/ng0197-74

5 Albrecht, U. et al. Imprinted expression of the murine Angelman syndrome gene, Ube3a, in hippocampal and Purkinje neurons. Nat Genet 17, 75–78 (1997). 10.1038/ng0997-75

6 Rougeulle, C., Glatt, H. & Lalande, M. The Angelman syndrome candidate gene, UBE3A/E6-AP, is imprinted in brain. Nat Genet 17, 14–15 (1997). 10.1038/ng0997-14

7 Copping, N. A., McTighe, S. M., Fink, K. D. & Silverman, J. L. Emerging Gene and Small Molecule Therapies for the Neurodevelopmental Disorder Angelman Syndrome. Neurotherapeutics 18, 1535–1547 (2021). 10.1007/s13311-021-01082-x

8 Elgersma, Y. & Sonzogni, M. UBE3A reinstatement as a disease-modifying therapy for Angelman syndrome. Dev Med Child Neurol 63, 802–807 (2021). 10.1111/dmcn.14831

9 Markati, T., Duis, J. & Servais, L. Therapies in preclinical and clinical development for Angelman syndrome. Expert Opin Investig Drugs 30, 709–720 (2021). 10.1080/13543784.2021.1939674

10 Meng, L. et al. Towards a therapy for Angelman syndrome by targeting a long non-coding RNA. Nature 518, 409–412 (2015). 10.1038/nature13975

11 Vendrame, M. et al. Analysis of EEG patterns and genotypes in patients with Angelman syndrome. Epilepsy Behav 23, 261–265 (2012). 10.1016/j.yebeh.2011.11.027

12 Valente, K. D. et al. Angelman syndrome: difficulties in EEG pattern recognition and possible misinterpretations. Epilepsia 44, 1051–1063 (2003). 10.1046/j.1528-1157.2003.66502.x

13 Levin, Y. et al. Evaluation of electroencephalography biomarkers for Angelman syndrome during overnight sleep. Autism Res 15, 1031–1042 (2022). 10.1002/aur.2709

14 Sidorov, M. S. et al. Delta rhythmicity is a reliable EEG biomarker in Angelman syndrome: a parallel mouse and human analysis. J Neurodev Disord 9, 17 (2017). 10.1186/s11689-017-9195-8

15 Frohlich, J. et al. Electrophysiological Phenotype in Angelman Syndrome Differs Between Genotypes. Biol Psychiatry 85, 752–759 (2019). 10.1016/j.biopsych.2019.01.008

16 Martinez, L. A. et al. Quantitative EEG Analysis in Angelman Syndrome: Candidate Method for Assessing Therapeutics. Clin EEG Neurosci 54, 203–212 (2023). 10.1177/1550059420973095

17 Ostrowski, L. M. et al. Delta power robustly predicts cognitive function in Angelman syndrome. Ann Clin Transl Neurol 8, 1433–1445 (2021). 10.1002/acn3.51385

18 Hipp, J. F., Frohlich, J., Keute, M., Tan, W. H. & Bird, L. M. Electrophysiological Abnormalities in Angelman Syndrome Correlate With Symptom Severity. Biol Psychiatry Glob Open Sci 1, 201–209 (2021). 10.1016/j.bpsgos.2021.05.003

19 Hipp, J. F. et al. The UBE3A-ATS antisense oligonucleotide rugonersen in children with Angelman syndrome: a phase 1 trial. Nature Medicine, 1–10 (2025).

20 Rotaru, D. C., Mientjes, E. J. & Elgersma, Y. Angelman Syndrome: From Mouse Models to Therapy. Neuroscience 445, 172–189 (2020). 10.1016/j.neuroscience.2020.02.017

21 Judson, M. C. et al. GABAergic Neuron-Specific Loss of Ube3a Causes Angelman Syndrome-Like EEG Abnormalities and Enhances Seizure Susceptibility. Neuron 90, 56–69 (2016). 10.1016/j.neuron.2016.02.040

22 Born, H. A. et al. Strain-dependence of the Angelman Syndrome phenotypes in Ube3a maternal deficiency mice. Sci Rep 7, 8451 (2017). 10.1038/s41598-017-08825-x

23 Born, H. A. et al. Early Developmental EEG and Seizure Phenotypes in a Full Gene Deletion of Ubiquitin Protein Ligase E3A Rat Model of Angelman Syndrome. eNeuro 8 (2021). 10.1523/ENEURO.0345-20.2020

24 Copping, N. A. & Silverman, J. L. Abnormal electrophysiological phenotypes and sleep deficits in a mouse model of Angelman Syndrome. Mol Autism 12, 9 (2021). 10.1186/s13229-021-00416-y

25 Gu, B. et al. Cannabidiol attenuates seizures and EEG abnormalities in Angelman syndrome model mice. J Clin Invest 129, 5462–5467 (2019). 10.1172/JCI130419

26 Spencer, E. R. et al. Longitudinal EEG model detects antisense oligonucleotide treatment effect and increased UBE3A in Angelman syndrome. Brain Commun 4, fcac106 (2022). 10.1093/braincomms/fcac106

27 Berg, E. L. et al. Insulin-like growth factor-2 does not improve behavioral deficits in mouse and rat models of Angelman Syndrome. Mol Autism 12, 59 (2021). 10.1186/s13229-021-00467-1

28 Lee, D. et al. Antisense oligonucleotide therapy rescues disturbed brain rhythms and sleep in juvenile and adult mouse models of Angelman syndrome. Elife 12 (2023). 10.7554/eLife.81892

29 Dickinson, A. H. et al. Atypical alpha oscillatory EEG dynamics in children with Angelman syndrome. NeuroImage: Clinical, 103865 (2025). 10.1016/j.nicl.2025.103865

30 Donoghue, T. A systematic review of aperiodic neural activity in clinical investigations. medRxiv, 2024.2010. 2014.24314925 (2024).

31 Donoghue, T. et al. Parameterizing neural power spectra into periodic and aperiodic components. Nat Neurosci 23, 1655–1665 (2020). 10.1038/s41593-020-00744-x

32 Wilkinson, C. L. et al. Developmental trajectories of EEG aperiodic and periodic components in children 2-44 months of age. Nat Commun 15, 5788 (2024). 10.1038/s41467-024-50204-4

33 Gentile, J. K. et al. A neurodevelopmental survey of Angelman syndrome with genotype-phenotype correlations. J Dev Behav Pediatr 31, 592–601 (2010). 10.1097/DBP.0b013e3181ee408e

34 Lossie, A. C. et al. Distinct phenotypes distinguish the molecular classes of Angelman syndrome. J Med Genet 38, 834–845 (2001). 10.1136/jmg.38.12.834

35 Moncla, A. et al. Angelman syndrome resulting from UBE3A mutations in 14 patients from eight families: clinical manifestations and genetic counselling. J Med Genet 36, 554–560 (1999).

36 Sonzogni, M. et al. A behavioral test battery for mouse models of Angelman syndrome: a powerful tool for testing drugs and novel Ube3a mutants. Mol Autism 9, 47 (2018). 10.1186/s13229-018-0231-7

37 Tanas, J. K. et al. Multidimensional analysis of behavior predicts genotype with high accuracy in a mouse model of Angelman syndrome. Transl Psychiatry 12, 426 (2022). 10.1038/s41398-022-02206-3

38 Boyd, S. G., Harden, A. & Patton, M. A. The EEG in early diagnosis of the Angelman (happy puppet) syndrome. Eur J Pediatr 147, 508–513 (1988). 10.1007/BF00441976

39 Bush, A., Zou, J. F., Lipski, W. J., Kokkinos, V. & Richardson, R. M. Aperiodic components of local field potentials reflect inherent differences between cortical and subcortical activity. Cereb Cortex 34 (2024). 10.1093/cercor/bhae186

40 Jeste, S. S., Frohlich, J. & Loo, S. K. Electrophysiological biomarkers of diagnosis and outcome in neurodevelopmental disorders. Current opinion in neurology 28, 110–116 (2015).

41 Negrón-Moreno, P. N., Diep, D. T., Guoynes, C. D. & Sidorov, M. S. Dissociating motor impairment from five-choice serial reaction time task performance in a mouse model of Angelman syndrome. Frontiers in Behavioral Neuroscience 16, 968159 (2022).

42 Sidorov, M. S. et al. Enhanced operant extinction and prefrontal excitability in a mouse model of Angelman syndrome. Journal of Neuroscience 38, 2671–2682 (2018).

43 Huang, H.-S. et al. Behavioral deficits in an Angelman syndrome model: effects of genetic background and age. Behavioural brain research 243, 79–90 (2013).

44 Van Woerden, G. M. et al. Rescue of neurological deficits in a mouse model for Angelman syndrome by reduction of αCaMKII inhibitory phosphorylation. Nature neuroscience 10, 280–282 (2007).

45 Godavarthi, S. K., Dey, P., Maheshwari, M. & Ranjan Jana, N. Defective glucocorticoid hormone receptor signaling leads to increased stress and anxiety in a mouse model of Angelman syndrome. Human Molecular Genetics 21, 1824–1834 (2012).

46 Senzai, Y., Fernandez-Ruiz, A. & Buzsaki, G. Layer-Specific Physiological Features and Interlaminar Interactions in the Primary Visual Cortex of the Mouse. Neuron 101, 500–513 e505 (2019). 10.1016/j.neuron.2018.12.009

47 Einstein, M. C., Polack, P. O., Tran, D. T. & Golshani, P. Visually Evoked 3-5 Hz Membrane Potential Oscillations Reduce the Responsiveness of Visual Cortex Neurons in Awake Behaving Mice. J Neurosci 37, 5084–5098 (2017). 10.1523/JNEUROSCI.3868-16.2017

48 Nestvogel, D. B. & McCormick, D. A. Visual thalamocortical mechanisms of waking state-dependent activity and alpha oscillations. Neuron 110, 120–138 e124 (2022). 10.1016/j.neuron.2021.10.005

49 Riyahi, P., Phillips, M. A., Boley, N. & Colonnese, M. T. Experience Dependence of Alpha Rhythms and Neural Dynamics in the Mouse Visual Cortex. J Neurosci 44 (2024). 10.1523/JNEUROSCI.2011-22.2024

50 McCormick, D. A. & Pape, H. C. Noradrenergic and serotonergic modulation of a hyperpolarization-activated cation current in thalamic relay neurones. J Physiol 431, 319–342 (1990). 10.1113/jphysiol.1990.sp018332

51 Soltesz, I. et al. Two inward currents and the transformation of low-frequency oscillations of rat and cat thalamocortical cells. J Physiol 441, 175–197 (1991). 10.1113/jphysiol.1991.sp018745

52 David, F. et al. Essential thalamic contribution to slow waves of natural sleep. J Neurosci 33, 19599–19610 (2013). 10.1523/JNEUROSCI.3169-13.2013

53 Lewis, L. D. et al. Thalamic reticular nucleus induces fast and local modulation of arousal state. Elife 4, e08760 (2015). 10.7554/eLife.08760

54 Amzica, F. & Steriade, M. Electrophysiological correlates of sleep delta waves. Electroencephalogr Clin Neurophysiol 107, 69–83 (1998). 10.1016/s0013-4694(98)00051-0

55 Moradi Chameh, H. et al. Diversity amongst human cortical pyramidal neurons revealed via their sag currents and frequency preferences. Nature communications 12, 2497 (2021).

56 Lendner, J. D. et al. An electrophysiological marker of arousal level in humans. Elife 9 (2020). 10.7554/eLife.55092

57 Gao, R., Peterson, E. J. & Voytek, B. Inferring synaptic excitation/inhibition balance from field potentials. Neuroimage 158, 70–78 (2017). 10.1016/j.neuroimage.2017.06.078

58 Wallace, M. L., Burette, A. C., Weinberg, R. J. & Philpot, B. D. Maternal loss of Ube3a produces an excitatory/inhibitory imbalance through neuron type-specific synaptic defects. Neuron 74, 793–800 (2012). 10.1016/j.neuron.2012.03.036

59 Wallace, M. L., van Woerden, G. M., Elgersma, Y., Smith, S. L. & Philpot, B. D. Ube3a loss increases excitability and blunts orientation tuning in the visual cortex of Angelman syndrome model mice. J Neurophysiol 118, 634–646 (2017). 10.1152/jn.00618.2016

60 Rotaru, D. C., van Woerden, G. M., Wallaard, I. & Elgersma, Y. Adult Ube3a Gene Reinstatement Restores the Electrophysiological Deficits of Prefrontal Cortex Layer 5 Neurons in a Mouse Model of Angelman Syndrome. J Neurosci 38, 8011–8030 (2018). 10.1523/JNEUROSCI.0083-18.2018

61 Kramer, M. A. & Chu, C. J. A General, Noise-Driven Mechanism for the 1/f-Like Behavior of Neural Field Spectra. Neural Comput 36, 1643–1668 (2024). 10.1162/neco_a_01682

62 Potter, S. N. et al. Linking Angelman and dup15q data for expanded research (LADDER) database: a model for advancing research, clinical guidance, and therapeutic development for rare conditions. Ther Adv Rare Dis 5, 26330040241254122 (2024). 10.1177/26330040241254122

63 Burnham, K. P. & Anderson, D. R. Multimodel inference: understanding AIC and BIC in model selection. Sociological methods & research 33, 261–304 (2004).

