## Supplemental Figures and Tables for "Periodic and aperiodic contributions to EEG delta power are translatable and complementary Angelman syndrome biomarkers"

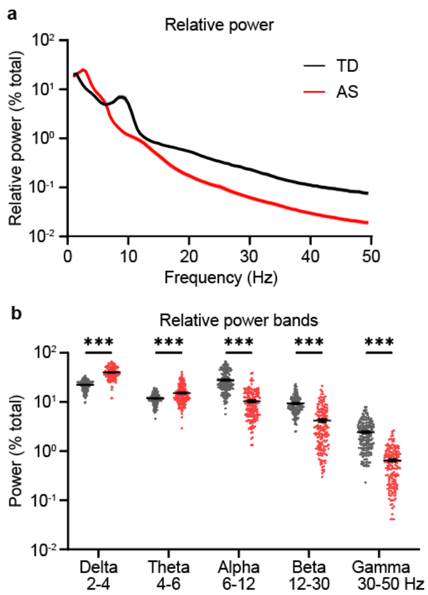


**Figure S1. Relative power analysis in TD and AS EEGs. a.** Relative PSDs from wake EEGs for all TD (black) and AS (red) EEGs. Relative power was calculated by normalizing raw PSD to total power for each EEG. Shading indicates ± SEM. **b.** Relative delta ($t(211.6) = 18.63$, $p < 0.001$) and theta ($t(208.7) = 5.90$, $p < 0.001$) power are increased in AS, and relative alpha ($t(276.9) = 15.62$, $p < 0.001$), beta ($t(335.4) = 12.25$, $p < 0.001$), and gamma ($t(228.4) = 14.27$, $p < 0.001$) power are decreased. TD: $n = 185$ recordings/subjects, AS: $n = 159$ recordings from 95 subjects. Points represent individual recordings, black lines indicate group averages ± SEM. *** $p < 0.001$, unpaired t-tests with Welch's correction, followed by Holm-Šídák correction for multiple comparisons.

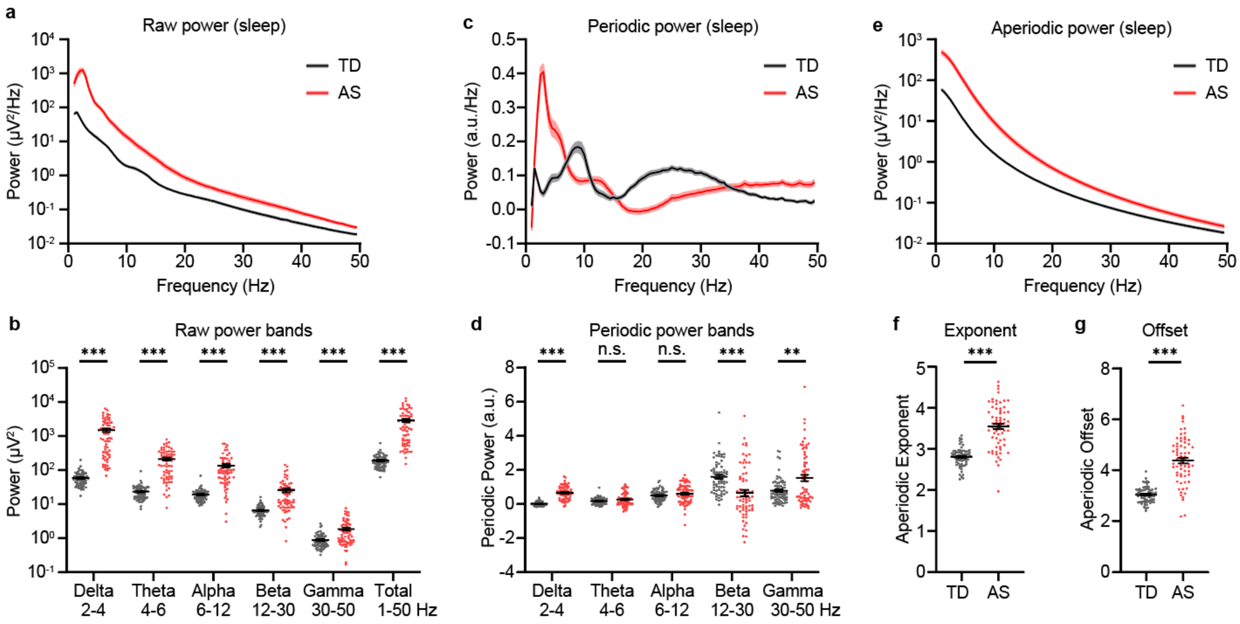


**Figure S2. Periodic and aperiodic power spectra reveal group differences between TD and AS EEGs during sleep. a.** Group PSDs from sleep EEGs (where available) for TD (black) and AS (red) EEGs. **b.** Raw power is elevated across all frequency bands for AS EEGs during sleep (delta: $t(67.04) = 7.88$, $p < 0.001$; theta: $t(67.57) = 8.74$, $p < 0.001$; alpha: $t(67.47) = 7.01$, $p < 0.001$; beta: $t(67.92) = 5.59$, $p < 0.001$; gamma: $t(76.46) = 4.84$, $p < 0.001$; total: $t(67.12) = 7.93$, $p < 0.001$). **c.** Group periodic power spectra calculated using *specparam*. **d.** Periodic delta ($t(77.25) = 13.19$, $p < 0.001$) and gamma ($t(92.96) = 3.60$, $p = 0.002$) power are increased in AS, and periodic beta ($t(112.1) = 4.36$, $p < 0.001$) power is decreased in AS. Periodic theta ($t(95.12) = 1.78$, $p = 0.15$) and alpha ($t(104.9) = 1.33$, $p = 0.19$) do not differ significantly by genotype. **e.** Group aperiodic spectra for TD and AS EEGs. **f.** Aperiodic exponent ($t(88.97) = 10.80$, $p < 0.001$) and **g.** offset ($t(80.80) = 11.67$, $p < 0.001$) are increased in AS EEGs. TD: $n = 71$ recordings/subjects, AS: $n = 68$ recordings from 56 subjects. **a,c,e:** Shading indicates ± SEM. **b,d,f,g:** Points represent individual recordings, black lines indicate group averages ± SEM. ** $p < 0.01$, *** $p < 0.001$, unpaired t-tests with Welch's correction, followed by Holm-Šídák correction for multiple comparisons.

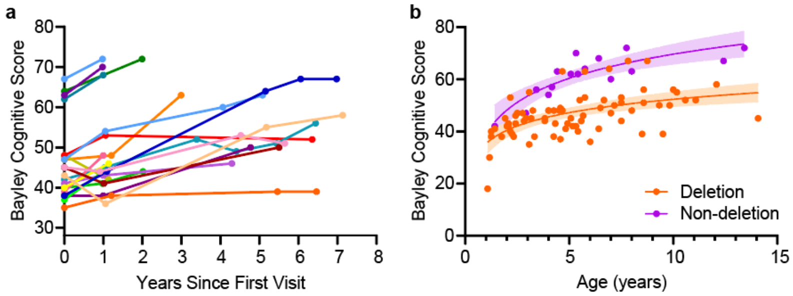


**Figure S3. Effects of age, genotype, and repeated measurements on Bayley cognitive scores. a.** Longitudinal changes in cognition among subjects with repeated measures (*n* = 19 subjects, 59 visits). Lines connect observations from the same individual. **b.** Effects of age and genotype on Bayley cognitive scores. Each point represents a single visit, colored by genotype (deletion: orange, *n* = 72, $R^2$ = 0.314; non-deletion: purple, *n* = 16, $R^2$ = 0.743). Lines show semi-log regression fits with shaded 95% confidence intervals.

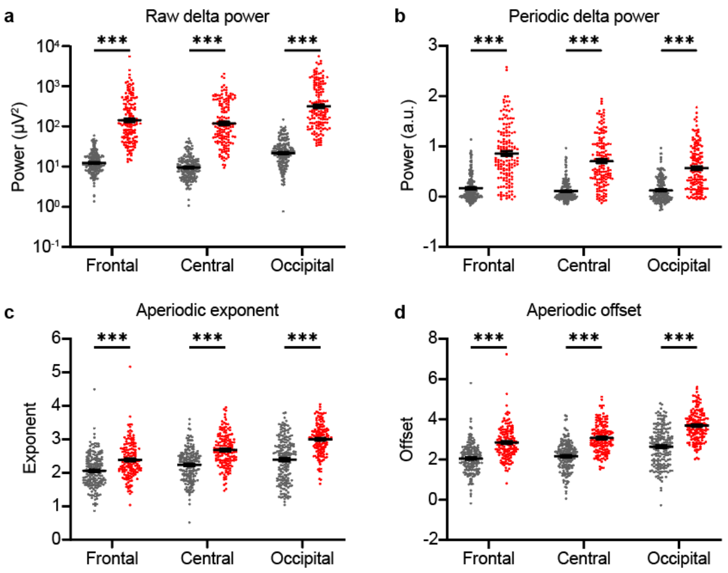


**Figure S4. Periodic and aperiodic AS phenotypes vary by cortical region.** Group differences for frontal, central, and occipital electrodes for different spectral features. Type-III ANOVA models were used to assess the effects of genotype and brain region on each EEG spectral feature, with subject modeled as a random effect and age included as a covariate. **a.** For raw delta power, there were significant main effects of genotype ($F(1, 333) = 246.01$, $p < 0.001$), region ($F(3, 1002) = 837.30$, $p < 0.001$), and a genotype × region interaction ($F(3, 1002) = 13.47$, $p < 0.001$). **b.** For periodic delta power, we found main effects of genotype ($F(1, 333) = 44.30$, $p < 0.001$), region ($F(3, 1002) = 102.16$, $p < 0.001$), and a strong genotype × region interaction ($F(3, 1002) = 56.80$, $p < 0.001$). **c.** Aperiodic exponent showed significant effects of genotype ($F(1, 333) = 167.16$, $p < 0.001$), region ($F(3, 1002) = 246.14$, $p < 0.001$), and genotype × region interaction ($F(3, 1002) = 21.63$, $p < 0.001$). **d.** Finally, for aperiodic offset, genotype ($F(1, 333) = 213.05$, $p < 0.001$), region ($F(3, 1002) = 298.83$, $p < 0.001$), and genotype × region interaction ($F(3, 1002) = 8.96$, $p < 0.001$) all reached statistical significance. **a-d:** Points represent individual recordings for TD (black) and AS (red) subjects. Black lines indicate group averages. *** $p < 0.001$, Tukey's multiple comparisons test. Effect sizes of post-hoc tests reported in **Table S1**.

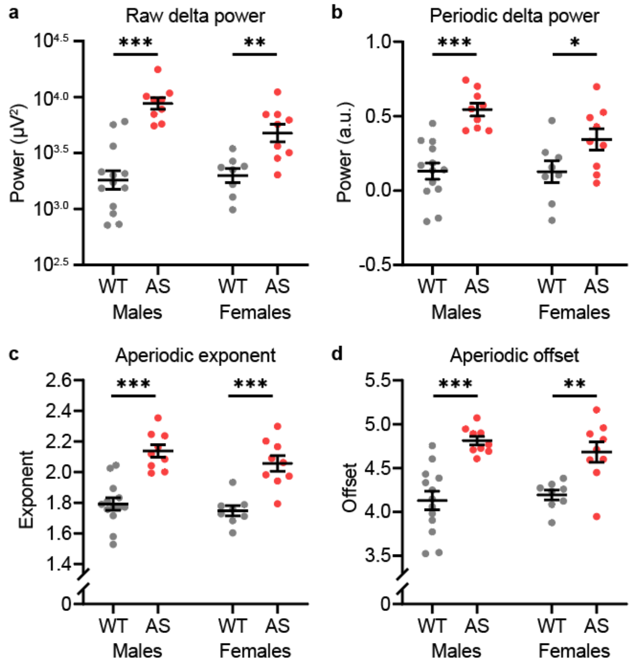


**Figure S5. Periodic and aperiodic phenotypes in mice separated by sex. a.** Raw delta power is elevated in both male and female Ube3a$^{m-/p+}$ mice (red) relative to WT (black). 2-way ANOVA: main effect of genotype: $F(1, 35) = 37.78$, $p < 0.001$; main effect of sex: $F(1, 35) = 2.11$, $p = 0.16$; genotype × sex interaction: $F(1, 35) = 3.90$, $p = 0.056$. **b.** Periodic delta is elevated in both male and female *Ube3a*$^{m-/p+}$ mice (genotype: $F(1, 35) = 39.04$, $p < 0.001$; sex: $F(1, 35) = 2.73$, $p = 0.11$; interaction: $F(1, 35) = 2.52$, $p = 0.12$). **c.** Aperiodic exponent is elevated in both male and female Ube3a$^{m-/p+}$ mice (genotype: $F(1, 35) = 56.57$, $p < 0.001$; sex: $F(1, 35) = 2.04$, $p = 0.16$; interaction: $F(1, 35) = 0.18$, $p = 0.67$). **d.** Aperiodic offset is elevated in both male and female *Ube3a*$^{m-/p+}$ mice (genotype: $F(1, 35) = 36.61$, $p < 0.001$; sex: $F(1, 35) = 0.12$, $p = 0.73$; interaction: $F(1, 35) = 1.01$, $p = 0.32$). **a-d.** Bars show mean ± SEM. Each point represents an individual mouse. Asterisks indicate results from unpaired t-tests with Welch's correction: *$p < 0.05$, **$p < 0.01$, ***$p < 0.001$.

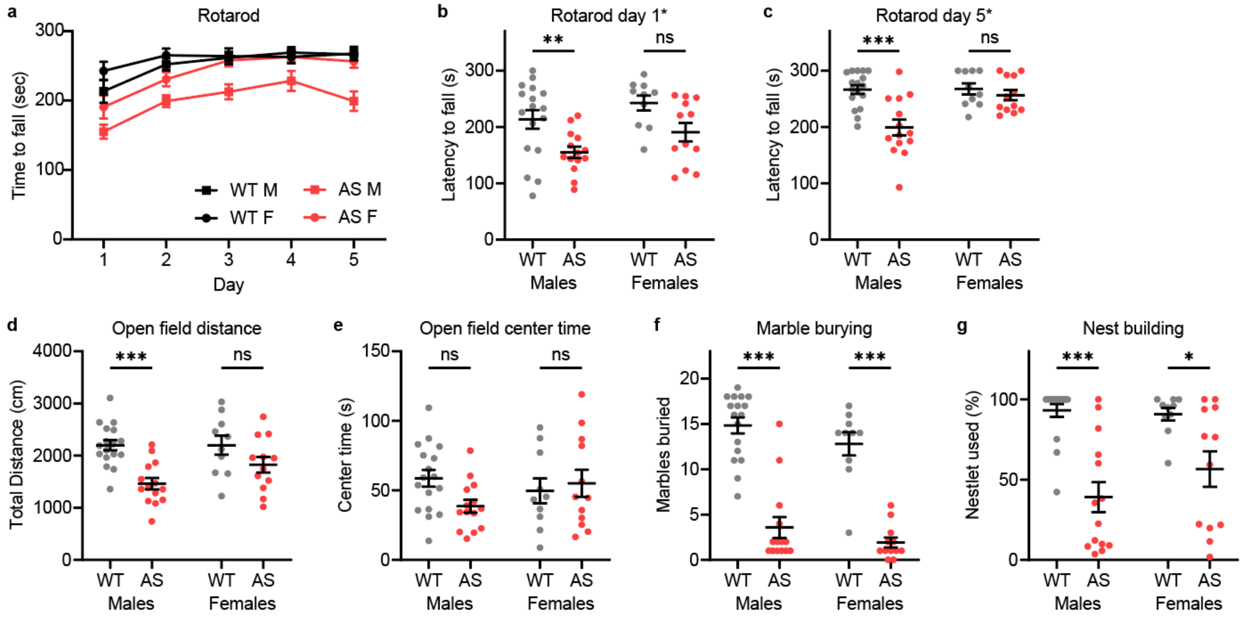


**Figure S6. Behavioral battery separated by sex.** Behavioral results with a significant main effect of sex or a sex × genotype interaction (indicated by asterisks on plot titles in **b-g**) were normalized separately by sex prior to PCA (Tanas, Kerr et al. 2022). **a.** Performance on rotarod across 5 days of testing, shown as mean latency to fall ± SEM. Black circles indicate WT females ($n$ = 8), black squares indicate WT males ($n$ = 13), red circles indicate $Ube3a^{m-/p+}$ females ($n$ = 11), and red squares indicate $Ube3a^{m-/p+}$ males ($n$ = 13). **b.** Day 1 latency to fall in rotarod task. 2-way ANOVA: main effect of genotype: $F(1, 49) = 13.30$, $p < 0.001$; main effect of sex: $F(1, 49) = 4.58$, $p = 0.037$; genotype × sex interaction: $F(1, 49) = 0.05$, $p = 0.83$. **c.** Day 5 latency to fall in rotarod task (genotype: $F(1, 49) = 13.42$, $p < 0.001$; sex: $F(1, 49) = 7.49$, $p = 0.009$; interaction: $F(1, 49) = 6.89$, $p = 0.012$). **d.** Total distance traveled in open field (genotype: $F(1, 49) = 17.88$, $p < 0.001$; sex: $F(1, 49) = 1.91$, $p = 0.17$; interaction: $F(1, 49) = 1.91$, $p = 0.17$). **e.** Time spent in center region of open field (genotype: $F(1, 49) = 1.00$, $p = 0.32$; sex: $F(1, 49) = 0.26$, $p = 0.61$; interaction: $F(1, 49) = 3.00$, $p = 0.090$). **f.** Number of marbles buried in marble burying task (genotype: $F(1, 49) = 120.3$, $p < 0.001$; sex: $F(1, 49) = 3.322$, $p = 0.075$; interaction: $F(1, 49) = 0.03$, $p = 0.86$). **g.** Percentage of nestlet used in nest building assay (genotype: $F(1, 49) = 32.24$, $p < 0.001$; sex: $F(1, 49) = 0.94$, $p = 0.34$; interaction: $F(1, 49) = 1.64$, $p = 0.21$). **b-g.** Each point represents an individual mouse. Black lines indicated mean ± SEM. Asterisks indicate results from Šídák's multiple comparisons test: *$p < 0.05$, **$p < 0.01$, ***$p < 0.001$.

| EEG Measure | Region | TD: mean ± *SD* | AS: mean ± *SD* | Effect size (Cohen's d) | |
|---|---|---|---|---|---|
| $Log_{10}$(raw delta power) (μV²) | Frontal | 1.10 ± *0.27* | 2.17 ± *0.55* | 2.53 | |
| | Central | 0.99 ± *0.28* | 2.09 ± *0.54* | 2.62 | |
| | Occipital | 1.35 ± *0.34* | 2.52 ± *0.55* | 2.61 | |
| | **Average** | **1.14 ± *0.28*** | **2.26 ± *0.54*** | **2.67** | |
| Periodic delta power (a.u.) | Frontal | 0.17 ± *0.25* | 0.86 ± *0.56* | 1.64 | |
| | Central | 0.11 ± *0.19* | 0.71 ± *0.50* | 1.62 | |
| | Occipital | 0.13 ± *0.24* | 0.57 ± *0.44* | 1.29 | |
| | **Average** | **0.14 ± *0.21*** | **0.71 ± *0.48*** | **1.61** | |
| Aperiodic exponent | Frontal | 2.06 ± *0.50* | 2.38 ± *0.53* | 0.62 | |
| | Central | 2.24 ± *0.48* | 2.67 ± *0.49* | 0.90 | * |
| | Occipital | 2.39 ± *0.66* | 2.99 ± *0.46* | 1.04 | |
| | **Average** | **2.23 ± *0.50*** | **2.68 ± *0.44*** | **0.95** | |
| Aperiodic offset | Frontal | 2.05 ± *0.73* | 2.83 ± *0.80* | 1.03 | |
| | Central | 2.16 ± *0.72* | 3.06 ± *0.72* | 1.25 | |
| | Occipital | 2.64 ± *0.96* | 3.69 ± *0.76* | 1.19 | |
| | **Average** | **2.28 ± *0.74*** | **3.20 ± *0.68*** | **1.27** | |

**Table S1. Regional comparisons of raw delta, periodic delta, and aperiodic features.** This table reports means ± SD and effect sizes (Cohen's d) for group differences between AS and TD EEGs. Frontal, central, and occipital regions correspond to F3/F4, C3/C4, and O1/O2 electrodes, respectively. All spectral features showed significant group differences across regions (see **Fig S4**). To evaluate regional variation in effect sizes, we compared region-specific *d* values using *z*-tests. For raw delta power, effect sizes were similar across regions (frontal vs. central: $z = 0.44$, $p = 0.95$; frontal vs. occipital: $z = 0.37$, $p = 0.95$; central vs. occipital: $z = -0.06$, $p = 0.95$). Periodic delta power showed a trend towards greater effect sizes at frontal and central sites (frontal vs. central: $z = -0.10$, $p = 0.92$; frontal vs. occipital: $z = -2.01$, $p = 0.084$; central vs. occipital: $z = -1.91$, $p = 0.084$). For aperiodic exponent, differences were more pronounced in occipital regions (frontal vs. central: $z = 1.76$, $p = 0.12$; frontal vs. occipital: $z = 2.62$, $p = 0.027$; central vs. occipital: $z = 0.86$, $p = 0.39$). Aperiodic offset did not show significant regional variation of effect sizes (frontal vs. central: $z = 1.32$, $p = 0.50$; frontal vs. occipital: $z = 0.97$, $p = 0.50$; central vs. occipital: $z = -0.35$, $p = 0.72$).

| | | PC1 | PC2 | PC3 | PC4 | PC5 | PC6 |
|---|---|---|---|---|---|---|---|
| **Loadings:** | Rotarod (day 1) | 0.395 | -0.321 | 0.442 | 0.648 | -0.152 | 0.320 |
| | Rotarod (day 5) | 0.346 | 0.188 | 0.704 | -0.354 | 0.047 | -0.471 |
| | Open field (distance) | 0.449 | 0.347 | -0.285 | 0.319 | 0.692 | -0.128 |
| | Open field (center time) | 0.244 | 0.746 | -0.113 | 0.063 | -0.570 | 0.205 |
| | Marbles buried | 0.518 | -0.218 | -0.127 | -0.590 | 0.096 | 0.558 |
| | Nest building | 0.442 | -0.370 | -0.446 | 0.031 | -0.402 | -0.553 |
| Total variance captured (%) | | 37.9 | 20.1 | 17.3 | 10.1 | 8.8 | 5.8 |
| Variance explained by genotype (%) | | 86.6 | 6.6 | 2.2 | 4.5 | 0.0 | 0.1 |

**Table S2. Principal component loadings and variance decomposition for analysis of mouse behavioral battery.** The table show loadings of six behavioral measures onto the first six principal components (PC1-PC6) from the principal component analysis shown in **Fig 5H**. Total variance captured indicates the proportion of total behavioral variance explained by each component. Variance explained by genotype reflects the proportion of variance in each component attributable to genotype, quantifying how well each PC separates *Ube3a*$^{m-/p+}$ and WT mice. Behavioral measures include rotarod performance (days 1 and 5), open field activity (total distance and center time), marble burying, and nest building.
